# Fish Metastasis Atlas for the Systematic Analysis of Organ-Specific Dissemination of Cancer

**DOI:** 10.64898/2026.11.10.765467

**Authors:** Hanieh Mazloom-Farsibaf, Ingrid Lekk, Qiongjing Zou, James F. Amatruda, Gaudenz Danuser

## Abstract

Metastasis is a multifaceted process in which cancer cells must overcome diverse biological barriers to establish secondary tumors and drive disease progression. Different cancers exhibit highly non-random, organ-specific patterns of dissemination, known as organotropism, which arise from cancer cell plasticity and interactions with the host microenvironment. A major challenge in the field is the lack of approaches that capture cancer cell dissemination at the whole-organism scale while maintaining physiological relevance. Here, we introduce an *in vivo* zebrafish model coupled with a data-driven computational framework to systematically map and compare patterns of cancer dissemination. Using patient-derived Ewing sarcoma cell lines, global analyses reveal pronounced heterogeneity across individual cell lines, which are further resolved by local metrics that capture region-specific colonization frequency and cancer cell visit density. This combined global–local approach identifies recurrent spatial preferences across the fish body and uncovers correlated organ targeting, indicating coordinated and structured dissemination dependent on cancer cell line and disease stage. The framework is robust and adaptable to diverse imaging datasets, providing a scalable platform for studying cancer– microenvironment interactions and spatially organized metastatic behavior for various cancer types as well as genetic and drug perturbations, with potential relevance for personalized treatment strategies.

## Introduction

Metastasis is the primary cause of cancer-related mortality^1,2^. During the metastatic cascade, cancer cells undergo a series of defined stages including local invasion, intravasation, survival in circulation, extravasation, and colonization at distant tissues. The early stages, which involve overcoming physical and biological barriers of the primary tumor tissue to dissemination, are generally similar across cancers. However, the ability of cancer cells to survive and proliferate within target tissue environments is highly context-dependent^3–5^. Indeed, metastatic spread is not random. Rather than disseminating indiscriminately, many tumors exhibit organotropism, a tendency to colonize specific organs^6^. For example, breast cancer commonly metastasizes the liver, lung, and bone^4,6–8^, whereas prostate cancer predominantly spreads to bone^9^. These patterns are not determined solely by cancer type; even cancer cells within a single tumor or across different patients with the same oncologic diagnosis exhibit high heterogeneity, leading to distinct organ-specific behaviors during metastasis^10,11^. Organotropism can arise from both genetic variation within tumor subpopulations^12,13^ and non-genetic plasticity^14,15^. Non-genetic reprogramming can reshape transcriptional, translational, and molecular networks, ultimately influencing how cancer cells interact with, and even remodel, the local microenvironment to support aggressive colonization^6^. Understanding organ-specific metastatic patterns provides insight into the mechanisms by which cancer cells adapt to distinct distant tissue niches, which is critical for developing targeted therapies for metastatic disease.

Ewing Sarcoma (EwS) is cancer in dire need of such investigations. EwS is an aggressive cancer of bone and soft tissue that primarily affects children, adolescents, and young adults. The survival rate of metastatic or recurrent Ewing sarcoma is <30% – an outcome that has not improved much in the past decades despite intensification of therapy and trials of experimental agents^16,17^. Furthermore, immunotherapies in pediatric solid tumors have not shown considerable success partly due to immunosuppressive tumor microenvironment (TME)^18^. Therefore, better understanding of the mechanisms of EwS metastasis, organotropism and colonization of new premetastatic niches in the TME is much needed to create effective targeted therapies. EwS arises from chromosomal translocations that fuse a member of the FET family of proteins to an ETS family transcription factor. The most common form of EwS (85% of cases) derives from a t(11;22)(q24;q12) translocation, which generates an oncogenic chimeric protein EWSR1::FLI1 (EF1). The mutational burden of EwS is otherwise low^14^. High levels of EF1 expression in EwS cells have been linked to increased proliferation^19,20^, whereas knock-down of EF1 leads to more protrusive cell morphology^19,21^ and increased spreading^19,20^, suggesting that in this case metastasis is possibly governed by a delicate balance between two competing functions.

Capturing the effects of these multifactorial and dynamic processes on the metastatic propensity of a cancer requires experimental systems that enable systematic, whole-organism tracking of cancer cell dissemination and organ-specific colonization. Several model systems have been developed to study organotropism, including mouse xenografts^22–24^, zebrafish models^25–27^, and human organoid platforms^28,29^. Mouse models provide physiological relevance but are limited in throughput and real-time visualization^30,31^. Human organoids offer tissue specificity but often lack systemic interactions such as vascular networks and physiological oxygen and nutrient gradients and therefore cannot fully recapitulate the multi-stage metastatic process occurring within an intact organism^32,33^.

In this study, we present the Fish Metastasis Atlas (FMA) framework to investigate the organotropic behavior of EwS cells using a zebrafish xenograft model. Zebrafish are uniquely suited for organism-level modeling of metastasis due to their capacity to produce hundreds of embryos in one pairing and the optical transparency of their larvae^31^. This makes zebrafish larvae ideal for systematic imaging and provides statistical power to study the heterogeneity of cancer cell spreading. Moreover, zebrafish share 70% orthologous genes with humans (85% when considering human disease-associated genes), and have a complex lymphatic, vascular, and immune system^34–36^. At the larval stage, the adaptive immune system of zebrafish has not yet developed, allowing xenografting experiments in immunocompetent wild-type animals. These advantages have rendered zebrafish a uniquely valuable model organism for cancer biology^34,36^.

Leveraging the features of the zebrafish, the proposed Fish Metastasis Atlas framework provides a quantitative, data-driven platform to study organ-specific spreading patterns of metastatic cancer cells *in vivo*. By using a reference image of vascular structures to register imaging data, it enables consistent alignment across individual animals and experimental conditions. This approach supports integration and expansion of datasets, facilitating cross-experimental comparisons, and the construction of cumulative atlases of metastatic dissemination. FMA allows systematic mapping of cancer cell distribution, identification of the top-ranked preferred regions, and comparative analysis across cell lines, tumor stages, and experimental perturbations. Additionally, the framework can be used to explore inter-regional relationships, providing unprecedented insight into coordinated patterns of site colonization across different conditions and cancer cell states.

## Results

### Designing Human Cancer Xenotransplantation in Zebrafish

Following a well-established method of human cancer-cell xenografting into zebrafish^37^ (Fig. 1a), we xenografted six different fluorescently labeled Ewing sarcoma (EwS) cell lines into the perivitelline space (PVS) at 2 days post fertilization (dpf) and imaged the larvae for metastatic spread at three days post injection (3 dpi). Of these EwS cell lines, TC32 and CHLA9 have been established from a primary tumor at diagnosis, whereas CHLA10 and CHLA258 come from a post chemotherapy metastatic lymph node and lung metastasis, respectively^38,39^. The comparison between CHLA9 and 10 is of special interest, as these cell lines were established from the same patient. Intermediate levels of EF1 have been correlated with higher metastatic potential^19,21,40^. We therefore used TC32 cells expressing doxycyline-inducible EF1 shRNA^21,41^, leading to decreased levels of EF1 expression (Supplementary Fig 1a) and allowing us to compare spread of TC32 with high (TC32 shEF1 CTR) and reduced (TC32 shEF1 KD) expression of the fusion oncogene EF1.

**Figure 1.**
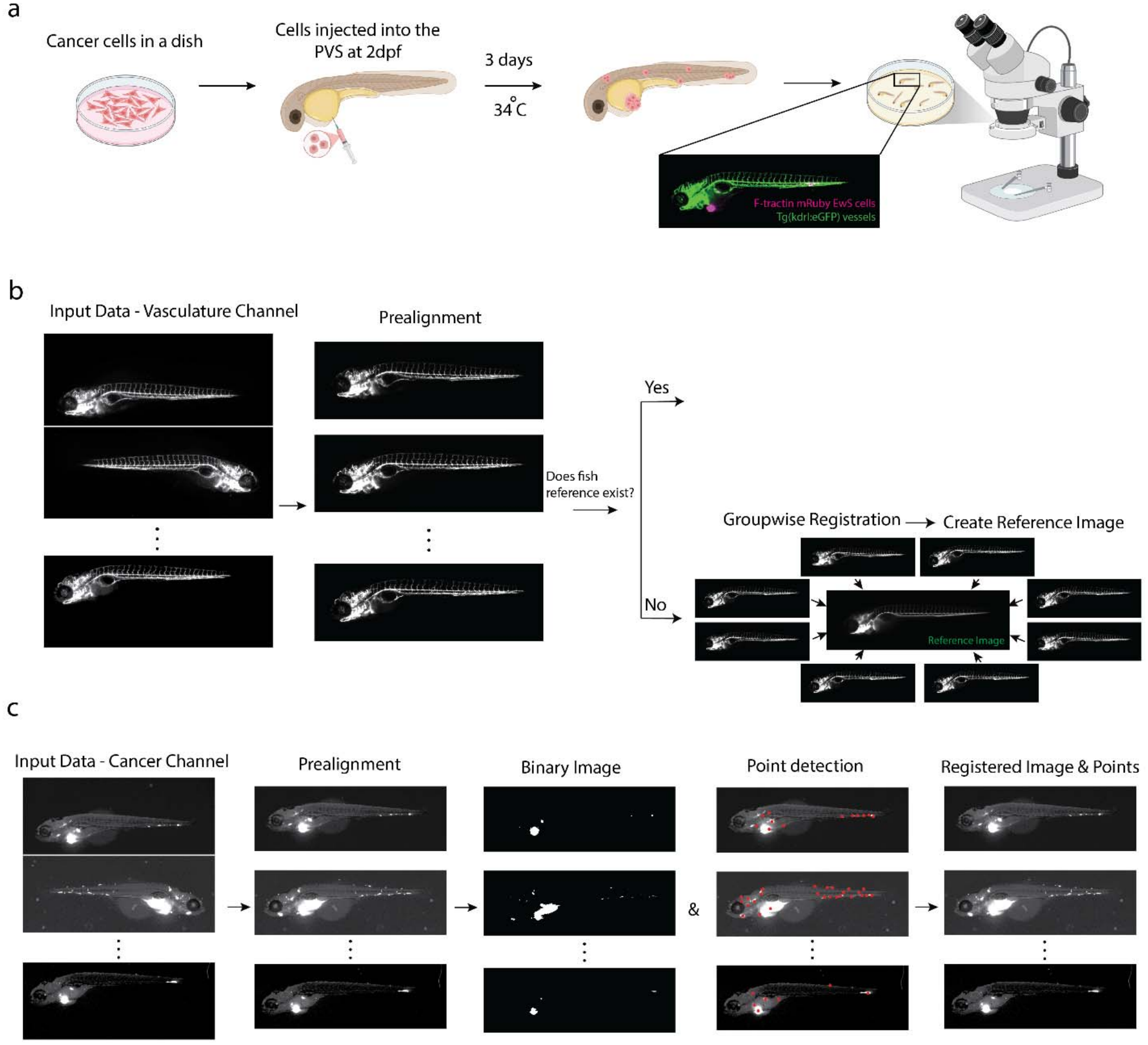
FishAtlas Workflow. a) Experimental workflow of the FishAtlas pipeline. Cancer cells were cultured *in vitro* and injected into the perivitelline space (PVS) of 2-day post fertilization (dpf) zebrafish larvae. At three days post-injection (dpi), larvae were imaged live by a two-channel dissection microscope: vasculature channel (*kdrl* transgenic line) and genetically labelled EwS cell channel. b) Registration workflow of fish images. Images were first pre-aligned. Depending on the availability of a fish reference, registration was performed using either pairwise or group-wise approaches, including rigid and deformable transformations. c) Cancer cell analysis workflow. Pre-aligned cancer cell images were registered with their corresponding vasculature images. Cancer colonies were detected at the individual fish level, producing both a binary mask representing colony locations and a point representation indicating each colony’s position for visitation frequency analysis. Each cancer image (binary or point representation) was then transformed using the registration map to generate accumulated visitation patterns across the population.

In the xenografting experiment, *Tg(kdrl:eGFP)* or *Tg(kdrl:dsRed)* fish were used to visualize the vasculature, whereas cells were labelled for a filamentous actin marker F-tractin (eGFP or mRuby)^21^. Per cell line, a total number of 250-300 embryos were injected to obtain ∼100 successfully imaged and analyzed larvae at 3 dpi. The remaining ∼200 larvae had to be excluded because they were dead or sick, had lost the engraftment by 3 dpi, or were not imaged properly (tilted, out of focus, without vasculature fluorescence). The viability rate (per cent of healthy larvae at 3 dpi), engraftment rate (per cent of healthy larvae with PVS tumor at 3 dpi) and metastasis rate (per cent of healthy larvae with cells outside of the PVS at 3 dpi) were similar for most EwS cell lines averaging at 84, 57 and 72%, respectively (Supplementary Fig. 1b). However, the viability and engraftment rates of larvae with TC32 shEF1 KD xenografts were significantly lower, possibly due to the toxicity of doxycycline treatment, requiring a higher number of total embryo injections.

Anesthetized larvae were mounted in 1% low-melting agarose and live-imaged with a Leica M205FA stereoscope in bright-field (not shown), vasculature and cancer cell channel (Supplementary Fig. 1c). This simple imaging setup provides high experimental throughput and easy reproducibility without specialized microscopy techniques. Provided that the larvae are mounted flat against the dish, the vasculature in the trunk and tail is stereotypically aligned and clearly visible due to the small size and translucency of zebrafish larvae. However, imaging a single focal plane does not allow good resolution in the head of the zebrafish larvae, which would require z-stack acquisition.

### Vasculature-guided registration of individual fish larvae onto a dataset-driven reference template

To probe the tendency of different cancer cell lines to spread in distinct patterns, we developed a computational pipeline referred to as Fish Metastasis Atlas (FMA) (Fig. 1b and c). To capture heterogeneity in cancer cell behavior during early metastasis, we registered the ∼100 larval fish images per condition into a reference (Table 1) defined by the zebrafish vasculature. The vasculature is a spatially rich, relatively stereotyped fiduciary across the entire larva (Supplementary Fig. 2), enabling accurate alignment of both global anatomy and fine internal structures, particularly in the trunk and tail. Moreover, it is closely linked to the metastatic behavior of the cancer cells^42^. Unlike a single-template reference, this population-based framework captures inter-sample variabilities including differences in larval developmental speed and imaging conditions. New cell lines and imaging conditions can be incorporated without redesigning the framework because the template is continuously updatable. Thus, this approach provides a robust framework for both current studies and future extensions.

**Table 1.** Sample size and point statistics.

|  | **CHLA9** | **CHLA 10** | **CHLA 258** | **TC32** | **TC32<br>shEF1 CTR** | **TC32<br>shEF1 KD** |
|---|---|---|---|---|---|---|
| Number of fish | 101 | 92 | 98 | 102 | 91 | 87 |
| Number of fish<br>after registration | 101 | 92 | 95 | 102 | 91 | 87 |
| Number of points | 765 | 1336 | 493 | 1111 | 678 | 928 |
| Average number of points | 7.57 | 14.52 | 5.13 | 10.9 | 7.45 | 10.67 |

Fish images were first pre-aligned with the anterior positioned on the left and the dorsal side facing upward, to reduce variability and facilitate high-throughput registration (Fig. 1b). Images were then aligned using rigid and deformable transformations, either by registering each image to a predefined reference shared across all conditions or by constructing the reference directly from the population when no initial reference was available. We compared two strategies: generating a single global reference from all conditions or generating condition-specific references. The global reference ensures consistent scaling across conditions, uses an optimized method to deform vasculature image at individual fish level, and avoids re-registration during condition-to-condition comparisons, but is less robust to the addition of new conditions. In contrast, condition-specific references register all fish images and align them independently across experimental conditions, and provide greater flexibility for future dataset expansion without requiring a shared reference. However, additional registration is needed for comparative analysis between conditions.

For both strategies, we first performed rigid registration to a fixed image randomly selected from the fish population, followed by a groupwise registration at the deformation step. The groupwise approach registers all images simultaneously using a deformation model to minimize intensity variance, under the constraint that the average deformation across the population is zero. This defines an unbiased mean reference frame without explicitly selecting a single reference image^43^. We found comparable registration performance for both strategies implemented on the current dataset (see next section). Based on these results, we selected the condition-specific reference strategy as our alignment method of choice.

### Validation of the registration pipeline

We validated the registration process using manually annotated markers on vasculature images (Supplementary Fig. 3a) and compared its sensitivity to (i) global versus condition-specific definition of the reference (Supplementary Fig. 3b, 3c), and (ii) the choice of reference image in the rigid registration step (Supplementary Fig. 4). To quantify registration accuracy, distances between markers in individual fish and the reference images were calculated before and after registration. As shown in Supplementary Fig 3, 4, after registration these distances were consistently below 40 pixels (∼ 90 µm), corresponding to approximately the space between adjacent intersegmental vessels (ISVs). Overall, the registration preserved global body shape and dorsal anatomical features, particularly within the trunk region. However, distance deviations varied across individual fish bodies, reflecting both experimental and biological heterogeneity in the dataset (Supplementary Fig. 5). Particularly, some variability in data acquisition and imaging conditions was observed, including differences in fish orientation around the DLAV (dorsal longitudinal anastomotic vessel) and dimmer vasculature signals in the tail (Supplementary Fig 5a, 5b), which resulted in higher registration uncertainty in this region. Biologically, larval fish even at the same developmental stage (5 dpf) exhibited distinct anatomical variation due to differences in developmental speed and imaging time; for example, the sixth ISV (intersegmental vessel) develops differently relative to the DA (dorsal aorta) junction across individual fish (Supplementary Fig. 5c). In addition, due to variation in the Tg(*kdrl:eGFP)* labeling at this developmental stage and differences in imaging angle, ISVs may be detected as one or two vessels (arterial and/or venous ISV), with occasional lymphatic sprouts also visible^44^ (Supplementary Fig. 5d). These results highlight the heterogeneity of the fish body and experimental conditions and emphasize the necessity of using a high-throughput approach that sufficiently compensates for sample heterogeneity in the study of organ-specific distribution of cancer cells. Qualitative validation, shown as overlays of vasculature images with xenografted cancer colony images, along with their corresponding binary masks and point-based representations, further confirmed accurate alignment across individual fish (Supplementary Movies 1–6).

### Integration of Cancer Colony Distribution of Individual Fish to visualize condition-specific organotropism

After successful alignment of all samples for a particular condition, the fluorescence signal of cancer cells was mapped into accumulators that visualized the colonization pattern. At the individual fish level, we constructed two complementary representations: (i) a binary representation obtained by thresholding fluorescence signals to identify cancer colonies, and (ii) a point-based representation capturing colony frequency in a region (Fig. 1c). Both representations were then transformed using the registration mappings obtained during alignment.

Using the accumulators, we were able to examine cancer colony spatial distributions between different experimental conditions (Fig. 2a). By visual inspection we noted that the six Ewing sarcoma (EwS) cell lines spread into the distinct colonization patterns aside from the generally dense signal at the injection site. For cell lines that exhibited wide-spread distribution (e.g. CHLA10 and TC32 shEF1 KD), colonization patterns imitated vasculature structures, indicating cancer cell proximity to vessels. Importantly, these distinct patterns could only be discerned because of the integration of colony positions across tens of fish. At the level of individual fish the heterogeneity of colonization precludes any form of assessment of systematic spreading, as exemplified by the CHLA9 cell line in Fig 2b. Accumulation of cancer cells across all fish in Fig. 2b revealed significant aggregation of CHLA9 cell line colonies between the dorsal aorta and the posterior cardinal vein (an area annotated as VDA – ventral wall of the dorsal aorta; see Supplementary Fig. 2 for anatomical annotations). However, the positions of colonies in three individual CHLA9-injected fish indicated in red, green, and blue varied widely (Fig 2b). Fish ID 1 (red) contributed primarily to the VDA, with a few cells in intersegmental vessels (ISVs) and dorsal longitudinal anastomotic vessel (DLAV). Fish ID 2 (green) contributed small colonies at the DLAV near the hindbrain, and the posterior caudal hematopoietic tissue (CHT) in the tail. Fish ID 3 (blue) contributed colonies mainly to the VDA near the yolk and at the distal CHT. These observations underscore the importance of a high-throughput experimental approach to perform robust organ-specific analyses of colonization patterns.

**Figure 2.**
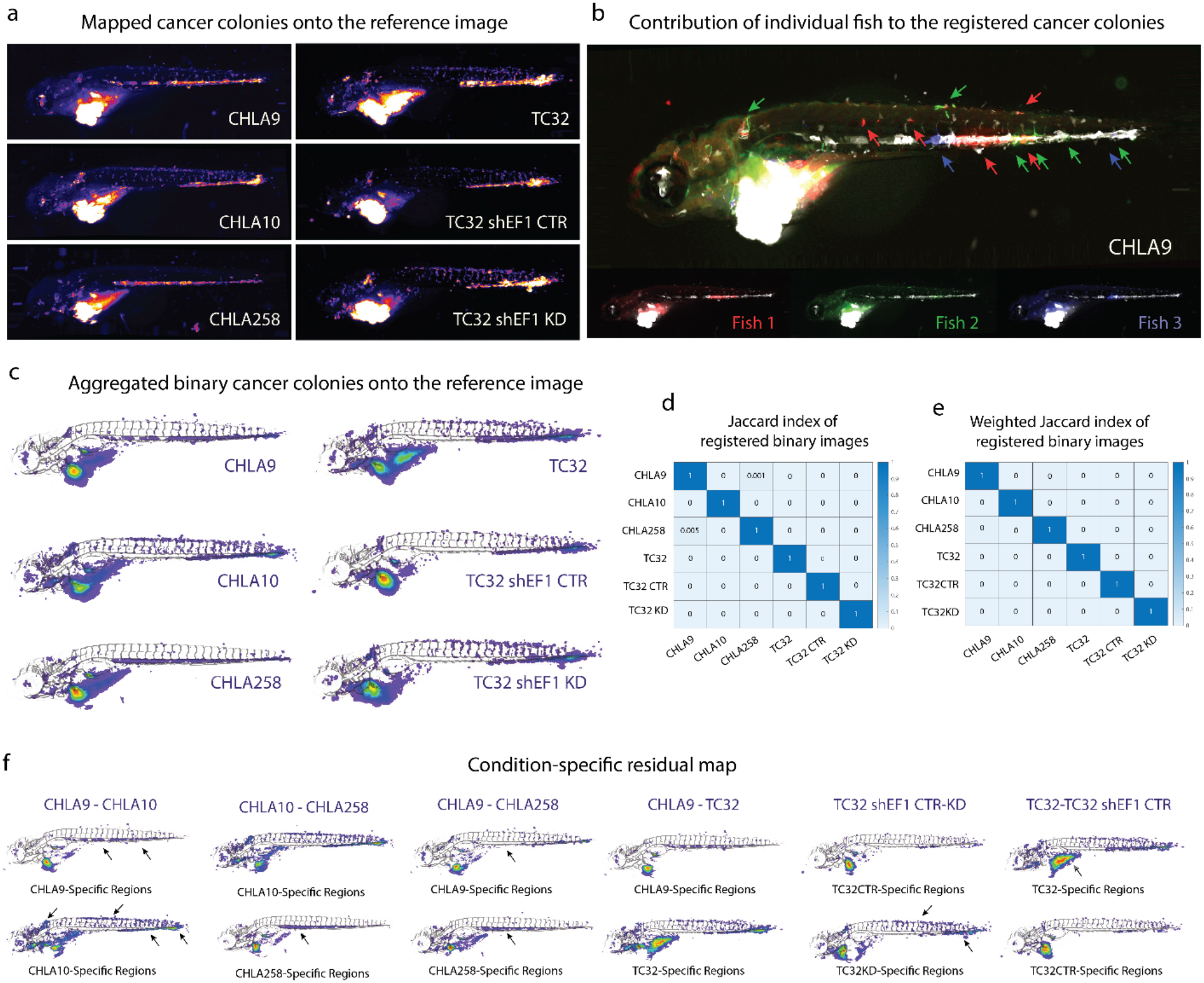
Accumulation of cancer cell colonization in a reference image. a) Accumulated maps of cancer colonization from six experimental conditions (Fire color) were registered to the same reference image. Maps are normalized by the number of fish per condition (n_CHLA9_ = 101, n_CHLA10_ = 92, n_CHLA258_ = 95, n_TC32_ = 101, n_TC32 shEF1 Ctr_ = 91, n_TC32 shEF1 KD_ = 87), such that a value of one (white) indicates colonization present in all fish for that condition. b) Three CHLA9-injected fish (shown in red, green, blue) are overlaid with the accumulated map of cancer colonization (n = 101 fish), all registered to a reference image. Insets of separate fish images illustrate the contributions of individual fish to the overall accumulation. c) Accumulated binary cancer colonization map across six experimental conditions, normalized to the number of fish per condition (rainbow color scale), and overlaid with the vasculature from the corresponding reference image. d) Similarity matrix of p-values summarizing pairwise comparisons among six conditions using the Jaccard index, calculated from accumulation maps thresholded at 1% of their maximum intensity. e) Similarity matrix of p-values summarizing pairwise comparisons among six conditions using the Jaccard index, calculated from grayscale accumulation maps, where pixel intensity reflects the number of fish. f) Six pairwise comparisons between cancer cell colonization patterns derived from residual maps as indicated. Residual maps display the first condition (e.g. CHLA9) after subtracting the overlapping regions with the second condition (e.g. CHLA10); that is, they show the regions unique to the first condition.

### Quantification of Differences in Colonization Patterns Across Ewing Sarcoma Cell Lines

Given the alignment of colonization patterns and vascular anatomy, we binarized colonization hotspots from individual fish, mapped them onto a vasculature reference image, and normalized them by fish number to compare occupied region profiles across conditions (Fig. 2c). Regions of consistently dense hotspots included the caudal hematopoietic tissue (CHT) and arteriovenous junction (AVJ) of the tail, branchial arches, eyes, jaw and the brain in the head, as well as intersegmental vessels (ISVs) and the dorsal longitudinal anastomotic vessel (DLA) in the trunk. Notably, some areas were colonized only by specific cell lines, such as CHLA9 and CHLA258 localization in the mid-trunk VDA.

To evaluate spatial similarity across conditions, we first measured pairwise Jaccard indices of the binarized accumulators (area of intersection over area of union), where any pixel containing at least one colony, including isolated cancer cells, was assigned a value of one. This analysis revealed similarity between CHLA9 and CHLA258, consistent with their visually similar mid trunk-dominant patterns (Fig 2d; shown are p-values of a statistical test with the null-hypothesis that the colonization patterns of two conditions are equivalent). To move beyond a binary representation of accumulation, we next computed Jaccard indices on grayscale accumulated maps, generated by integrating binary maps across individual fish such that grayscale intensity reflects the number of fish with cancer colonies at a given pixel (see Methods). Contrary to the binary analysis, the comparison of grayscale representations indicated that also the distributions of CHLA9 and CHLA258 differed significantly because of differences in the occupancy of the mid trunk region. Together, these results quantitatively confirm that each condition exhibits a distinct spatial colonization pattern.

### Identification and Analysis of Cell-Line–Specific Colonization Regions

We identified regions uniquely enriched for specific cell lines by performing pairwise comparisons across the six conditions after normalization to the different sample sizes (Fig. 2f). Comparing CHLA9 and CHLA10, derived from the same patient at diagnosis and from a metastatic lymph node, respectively, revealed distinct region-specific enrichment after excluding overlapped regions primarily at the injection sites. CHLA9 preferentially occupied regions proximal to the injection site, such as the VDA and anterior CHT (black arrows, Fig. 2f). CHLA10 showed enrichment also in the distal parts of the body, such as the caudal CHT, distal tail (AVJ), dorsal DLAV, ISVs (most markedly the most anterior ISV near the hindbrain) and the head. These data indicate that the metastatic cell line (CHLA10) exhibits increased spreading and colonization of new niches compared to the cell line originating from the primary tumor (CHLA9). We next compared two metastatic cell lines from different patients, CHLA10 and CHLA258, originating from a metastatic lymph node and lung metastasis, respectively. Surprisingly, CHLA258 exhibited a highly confined distribution in the VDA, compared to the wide-spread localization of CHLA10 colonies, perhaps highlighting the differences between cell-lines from different metastatic sites. In fact, as indicated already by the grayscale Jaccard analysis, the CHLA258 distribution was more similar to CHLA9 than CHLA10 (Fig 2c). Yet, after subtracting shared regions, it became obvious that the VDA was predominantly occupied by CHLA258 rather than CHLA9 (Fig 2f). Additionally, CHLA258 lacked the low-population colonies or single-cell spreading observed in ISVs and DLAV for CHLA9.

Both TC32 and CHLA9 are primary tumor–derived cell lines; however, their colonization patterns were largely exclusive, with overlap primarily restricted to the hindbrain. Outside this region, the spatial distributions resembled their original individual patterns, indicating minimal shared colonization regions. The distribution of TC32 overall resembled that of CHLA10. This was initially surprising, as TC32 is typically classified as a primary tumor-derived line, while CHLA10 derives from a metastatic lesion. This finding prompted us to look more closely at the original case report describing the clinical characteristics of the patient whose tumor gave rise to the TC32 line^45^. The patient was a 17 yo female who presented at an advanced stage of disease with a left ilium soft tissue mass and widespread metastasis to bones and bone marrow. Thus, biopsy of the tumor mass likely contained cells that had already acquired metastatic propensity, a feature that is recapitulated in the TC32 FMA data.

To look more specifically at the effect of EF1 expression level on EwS cell distribution, we took advantage of a TC32 derivative cell line incorporating an inducible shRNA specifically targeting EF1 (TC32-shEF1)^41^. We first established the colonization pattern of TC32-shEF1 cells at baseline (TC32-shEF1 CTR). Compared to the unmanipulated parental TC32 cell line, TC32 shEF1 CTR cells exhibited less colonization of distal sites such as the DLAV and ISV regions. This difference between two TC32 control conditions could reflect the genetic manipulation and selection that TC32 shEF1 CTR has undergone, together with variability in xenografting efficiency between the two cell lines, with TC32 shEF1 CTR forming smaller tumor masses at the injection site than TC32. Doxycycline-mediated EF1 knock-down (TC32 shEF1 KD) led to enrichment of xenografted cells toward the distal areas, with increased single-cell spreading along the DLAV and ISVs in the tail relative to the trunk. The TC32 shEF1 CTR residual map, obtained by subtraction from TC32 shEF1 KD, shows near-zero values, indicating that EF1 knock-down primarily enhances distal and low-density colonization rather than altering global distribution patterns. These results support the previously published data that lower levels of EF1 lead to higher metastatic potential^16,17^.

### Anatomical visitation frequencies of cancer cells

To complement the binary representation, which captures the spatial extent and size of colonization, we introduced a point-based representation in which each cancer colony is described as a single point, enabling quantification of how frequently cancer cells visit specific anatomical sites within the zebrafish body (Fig. 3a). Binary and point representations reveal broadly similar patterns across cell lines, while the point-based representation better preserves local and rare colonization events, including single-cell spreading that may be missed during binary map generation. Even when such events are retained after binarization, pixel-level transformation maps used during registration to a single reference image can distort or eliminate single-cell signals. By detecting colonies at the individual fish level and transforming their coordinates rather than pixelated regions, rare and single-cell events are retained.

**Figure 3.**
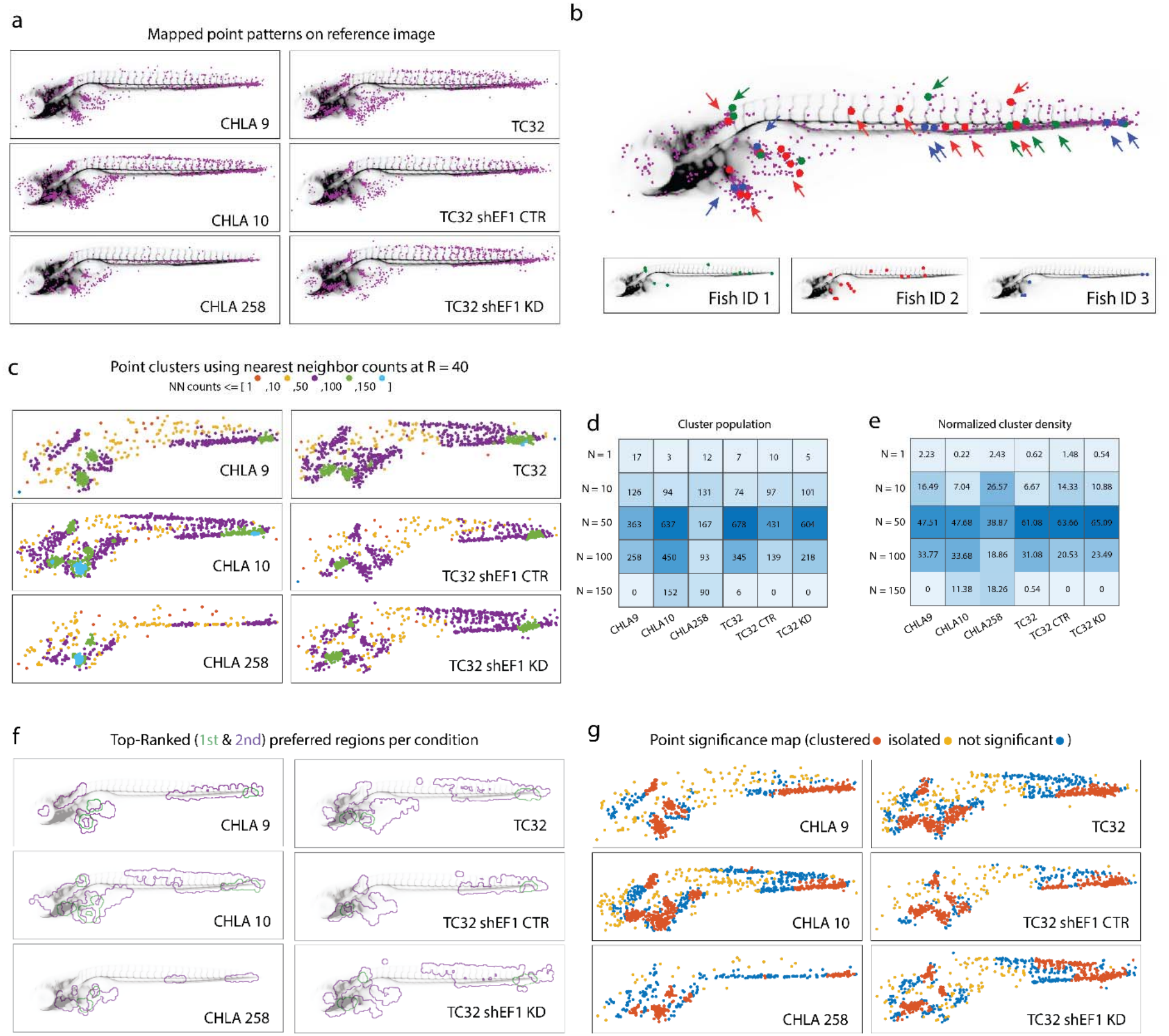
Cancer visitation patterns in the reference image. a) Accumulator patterns of cancer colony visitations from six experimental conditions in EWS cell–injected fish, all registered to the same reference image. The number of fish per condition is n = [101, 92, 95, 101, 91, 87], arranged from top to bottom, left to right. b) Three CHLA9-injected fish (red, green, blue) are shown together with the accumulated map of cancer colony visitation sites (n = 101 fish), all registered to a vasculature-guided reference image. Insets show individual contributions. c) Clustered point patterns based on the number of nearest neighbors within a radius of *r* = 40 pixels. Colors represent local density: red (1 neighbor), yellow (2–10), purple (11– 50), green (51–100), and cyan (101–150). d) Matrix indicating the absolute number of clusters in size bins across six experimental conditions. e) Matrix indicating the percentage of points falling with each of the size bins across six experimental conditions. f) Preferred regions for each experimental condition are identified by visitation frequency. Regions are ranked by local density, with the top two outlines in green and purple, based on the density class in (c). g) Classification of points as clustered or isolated based on nearest-neighbor analysis. Red points indicate significantly higher local neighbor counts than expected under random distribution (enrichment); yellow points indicate significantly lower neighbor counts (exclusion); and blue points indicate no significant difference from random distribution (using a permutation test, see Methods).

We quantified the total number of detected points and the average number of points per fish for each condition, revealing the propensity of each cell line to spread throughout the fish body; for example, CHLA10 exhibits approximately twice the spatial spread compared to CHLA9 (Table 1). We also quantified visitation frequency at specific anatomical sites. For instance, in CHLA9, binary maps show uniform occupancy of the CHT across the trunk and tail, while point patterns reveal higher visitation frequency in the tail than the trunk. Similarly, in CHLA258, aggregation within the VDA shows lower visitation in the trunk compared to the tail, a difference that is more clearly resolved in the point-based representation than in the binary maps. In CHLA10, some rare single-cell colonies along dorsal regions are missed in binary maps but appear in the point-based representation. Similar effects are observed across TC32 conditions, where rare dorsal colonization events are consistently underrepresented in binary maps. Overall, while binary maps indicate occupied regions, point patterns more accurately reflect visitation frequency and rare dissemination events throughout the fish body.

Individual fish contributed unevenly to organ-specific frequency patterns, as illustrated for CHLA9 (Fig. 3b): Fish ID 1 (red) and Fish ID 2 (green) contributed to the hindbrain and the mid-tail region (CHT), whereas Fish ID 3 (blue) primarily contributed to the mid-trunk and distal tail (AVJ). Fish-specific differences were also observed along distal vascular structures, with some fish contributing points along both the intersegmental vessels (ISVs) and the dorsal longitudinal anastomotic vessel (DLAV), while others contributed predominantly to the DLAV. These results highlight that integrating data from large sample populations is essential for meaningful analysis of organ-specific metastatic patterns.

### Density-based characterization of spatial colonization patterns

To further characterize spatial point patterns for individual cell lines, we quantified the local point density by counting for each colonization event the neighboring points within the radius of 40 pixels (∼ 90 µm) set by the registration accuracy (see above). Points were grouped into five density classes (N = 1, 10, 50, 100, 150) and visualized using distinct colors with edge correction applied for points near the fish body boundary (Fig 3c). This analysis revealed a continuum of spatial behaviors, ranging from sparse single-cell events to highly recurrent colonization sites. For each cell line, the distribution of cluster populations and per-fish density across these classes is summarized, providing a quantitative description of anatomical site-specific visitation patterns (Fig 3d, e).

Using local point density, we identified the top-ranked preferred anatomical regions for each cell line using an equal number of fish across conditions (n = 85) (Fig 3f). After excluding injection sites, which consistently ranked highest across conditions, the first two highest-ranked regions (green and purple outlines) differed markedly in size and/or location. CHLA9 preferentially targeted the ventral mid-trunk region; CHLA10 showed enrichment along the dorsal trunk and distal tail; and CHLA258 localized to multiple narrow regions within the ventral trunk without a clearly defined top-ranked region (green outlines), indicating a lower propensity for spreading compared with the other conditions, consistent with Table 1. The expression level of EF1 determined condition-dependent preferred regions, with broader spatial distributions in TC32 shEF1 KD compared with the more confined pattern observed in TC32 shEF1 CTR. By overlaying these regions onto the vasculature-guided reference image, we could also visualize the ranked regions within the context of vascular architecture (Supplementary Fig. SI6).

Finally, we compared each condition to randomly distributed point patterns containing the same number of points. The points were distributed across the fish body, defined by a body mask generated across all conditions. This analysis enabled identification of regions with fewer neighbors than expected (spatially isolated; yellow), and regions with an excess of neighbors (spatially clustered; red) from those that randomly distributed (blue), within a 40-pixel (∼ 90 µm) radius (Fig 3g). While both CHLA10 and TC32 shEF1 KD showed increased spreading to dorsal regions compared to TC32 shEF1 CTR, TC32 cells knocked down for EF1 expression exhibited higher local neighbor counts (red dots), suggesting that these cells may have a more stringent requirement for local environmental conditions to support their successful metastasis.

### Enriched anatomical regions

To complement automated identification of enriched regions based on local point density, we analyzed cell localization within user-defined anatomical regions of the reference fish. Eight biologically annotated regions were manually delineated (Fig 4a). We quantified the number of events per annotated region, normalized per fish (Fig. 4b). This analysis reflects the propensity of cancer cells across conditions spreading within each region. Specifically, vasculature regions, including the intersegmental vessels (ISVs) and the DLAV showed higher event frequencies in CHLA10 and TC32 shEF1 KD. In contrast, CHLA9 and CHLA258 exhibited increased localization in the VDA, and lower contributions to the branchial arches and jaw region compared to other conditions.

**Figure 4.**
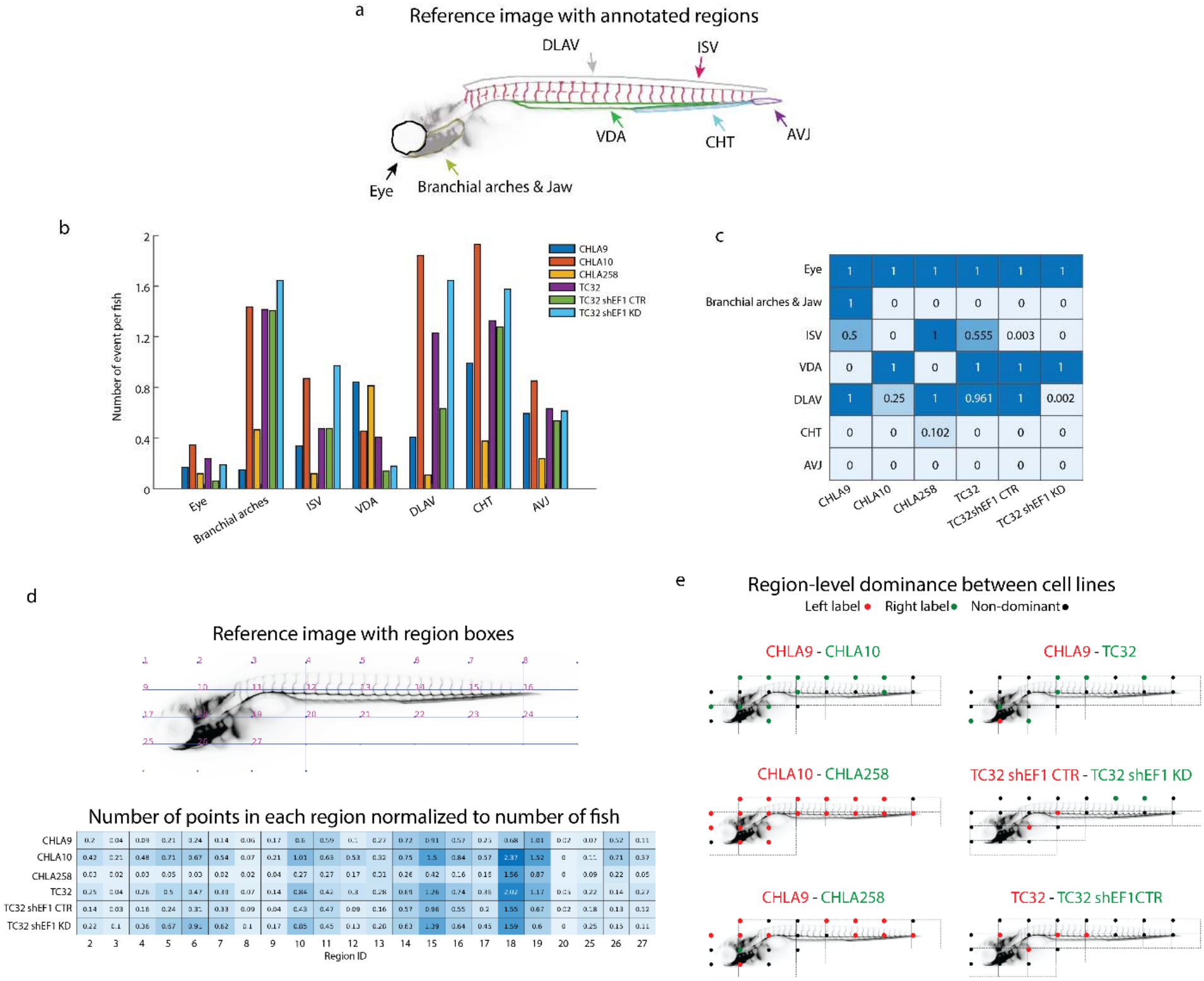
Region-based analysis of cancer colony visitation patterns in a reference image. a) Reference image of a fish with manually annotated regions outlining the eye, branchial arches and jaw, dorsal longitudinal anastomotic vessel (DLAV), intersegmental vessel (ISV), ventral wall of the dorsal aorta (VDA) (aorta-gonad mesonephros at earlier stages of development), caudal hematopoietic tissue (CHT), and atrioventricular junction (AVJ). b) Cancer colony visits per fish in each region. Colors represent different experimental conditions. c) Heatmap of p-values indicating whether a region is significantly enriched compared with a random distribution for each experimental condition. d) Box grid overlaid on the reference image, with a box size of [100 × 200] pixels. Distribution of points per region, normalized by the number of fish, representing the average frequency of region visitation per fish. e) Region-level dominance maps for six pairwise comparisons of experimental conditions. Red indicates dominance of the left-labeled condition; green indicates dominance of the right-labeled condition, and black indicates regions with no significant difference based on the Wilcoxon rank-sum test.

To statistically assess the enrichment, we computed *p*-values of a test under the null hypothesis a particular region is visited randomly, i.e. low p-values indicate strong preference for a cell line to spread to the analyzed region (Fig 4c). Enrichment in the caudal hematopoietic tissue (CHT) and the arteriovenous junction (AVJ) is consistent across all conditions except CHLA258. In contrast, enrichment in the ventral wall of the dorsal aorta (VDA), specifically the region between DA and PCV, is significant for CHLA9 and CHLA258. For the ISV region, enrichment is not significant in this analysis because the ISVs are defined across the entire fish body, including both trunk and tail. However, enrichment is high in the tail region for both TC32 shEF1 KD and CHLA10. In the DLAV, TC32 shEF1 KD exhibited additional enrichment that was not observed in CHLA10, consistent with previous results. These results also highlight that not all visually apparent clusters correspond to statistical enrichment: the eye region did not show significant enrichment in any condition, indicating that its occupancy was not distinguishable from random expectation, consistent with previous results based on local density analysis (Figure 3g). Overall, manual annotation of specific regions helps identify localized enrichment that is not fully captured by the previous analysis.

### Automated region-based analysis of colonization patterns

Finally, we also quantified point-based colonization events within a grid of sampling boxes across the reference image to identify region-specific patterns of colonization in an unbiased manner. In each box we measured the point aggregation and normalized it for variation in number of fish (Fig. 4d). Comparing colonization frequencies across regions for individual conditions (single row in Fig. 4) revealed patterns consistent with earlier enrichment analyses, with minor differences arising from region definitions that depend on box size.

Certain regions emerged as dominant anatomical hotspots across all conditions, including the injection site (box IDs 11, 19, 27 and part of 18) and the CHT and distal tail (boxes 14–16) (Fig. 4d). Some boxes included multiple anatomical regions, which are less reliable but still informative. For example, box 10 is a hotspot across all conditions, and captures part of the primordial channel and central arteries of the hindbrain as well as a few rostral ISVs. Additionally, certain regions were condition-specific, such as (i) the DLAV and tail ISVs (boxes 5–7) for CHLA10 and TC32 shEF1 KD; (ii) the hindbrain/first ISV (box 2) for CHLA9 and CHLA10; and (iii) the VDA between dorsal aorta and posterior cardinal vein (box 12-13) for CHLA258.

Considering the absolute values of normalized colonization, Fig 4d underlines again that CHLA10 disseminates approximately twofold stronger across enriched regions compared to CHLA9, indicating that the zebrafish model preserves the functional differences between metastatic and primary tumor in patient-matched cancer cell lines. TC32 shEF1 KD, and CHLA10 also shared elevated occupancy in dorsal parts of the tail (box 6), suggesting overlapping dissemination routes among these cell lines (Fig 4d).

To directly compare regional colonization between cell lines, we performed six pairwise comparisons (similar to Fig. 2f) using a nonparametric rank-based test (Fig. 4e). For each comparison, regions were classified according to which cell line showed dominant enrichment. This analysis highlighted regions in which one cell line consistently exhibited higher colonization frequencies over the other. In contrast, several regions did not differ significantly between conditions and are indicated as non-dominant (black circles) in the figure.

### Coordinated interdependencies between colonized regions

As the analyses thus far demonstrated organotropic behavior through regional enrichment, we wondered whether for certain cell lines the colonization patterns would exhibit structured interdependencies between anatomical regions. That is, if a particular cell line tends to colonize a certain anatomical region, would it then also colonize another region, possibly of similar environmental properties? To identify such relationships, we leveraged the sizable pool of injected fish per cell line and computed region-wise coexistence percentages based on fish-level event counts across five box-grid sizes chosen as multiples of 50 pixels (109 µm) (Supplementary Fig 7a, see Methods). Using the 100 × 200 grid as an example, regional coexistence patterns for the CHLA10 condition are shown in Figure 5a. Fish-level event counts for representative regional pairs with high and low coexistence are shown in Figure 5b, compared to a pair from randomly distributed pattern (left to right). Of note, co-occurrence analyses were restricted to non-zero values to avoid artificial inflation by shared absences across regions (see Methods).

**Figure 5.**
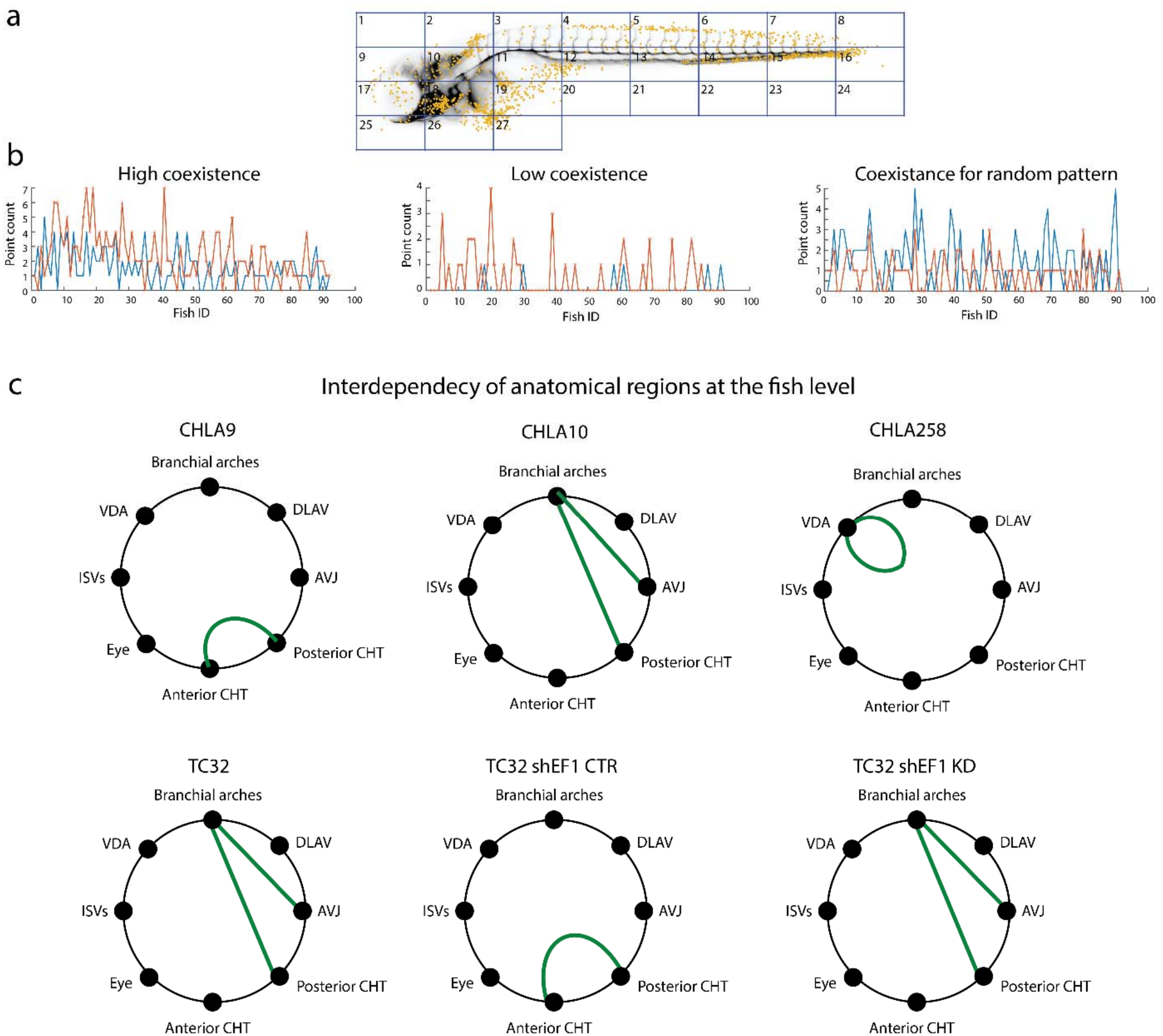
Interdependency of cancer colonies across fish body. a) Point pattern of CHLA10 cells registered to the reference image and overlaid with a box grid size of 100 × 200 pixels. b) Number of events per fish in selected two regions shown in blue and red color: (left), a high coexistence pair (regions 8 and 4); (middle), a low coexistence pair (regions 19 and 18); and (right), randomly distributed point patterns across the entire fish body, with two regions selected at random. c) Interdependencies between anatomical regions in visitations per fish. For each condition, nodes in a circular layout represent labeled anatomical regions. Green edges denote co-existence between regions, with edge thickness proportional to the magnitude of co-occurrence.

Examination of the maximum coexistence values revealed that very small grid sizes inflated coexistence due to sparse occupancy and noise sensitivity, whereas larger grids reduced spatial specificity by aggregating independent colonization events (Supplementary Fig 7b). Accordingly, an intermediate grid size of 100 x 100 pixels was selected to balance robustness and spatial resolution. Because the fish body is elongated along the anterior–posterior axis, rectangular grids were also evaluated to better align with its anatomical geometry. Specifically, grid sizes of 100 × 200 and 50 × 100 pixels were tested to assess the consistency of regional coexistence patterns across different box sizes.

Across conditions, regional pairs exhibiting high coexistence were consistently identified across grid sizes after excluding regions near the injection site and swim bladder. Coexistence magnitudes depended on experimental condition and box size, as the total number of points varied across conditions, and coexistence was sensitive to both grid size and sampling density. Accordingly, conditions with higher numbers of points, such as CHLA10, TC32, and TC32 shEF1 KD (see Table 1) generally exhibited higher coexistence values. For visualization, a fixed coexistence threshold of 0.45 was applied across all box sizes and conditions, corresponding to regional pairs in which at least 45% of events co-occurred at the single fish level (Supplementary Fig 8a). However, a more rigorous approach for evaluating co-occurrence is to define a threshold specific to each condition and box size (Supplementary Fig 8b).

To address this, we generated 100 randomly distributed point patterns per condition to compute condition-specific thresholds. The significance threshold was defined as the mean plus one standard deviation of the maximum coexistence values obtained from these random distributions (Supplementary Fig. 9). We then retained regional pairs that were consistent across spatial scales (Table 2). Beyond the significance of threshold, an additional consideration is the prevalence of regional coexistence across fish. While this framework identifies whether regional coexistence occurs, it does not quantify how frequently it occurs across the population. We therefore report the number (fraction) of fish exhibiting regional coexistence for each pair (Table 2).

**Table 2.** Jaccard index for regional coexistence across box sizes. C denotes the threshold derived from random distributions, calculated as the mean plus standard deviation (C = *μ_n_*_=100_ + *σ_n_*_=100_) of the maximum Jaccard indices between regions.

|  | **[50 100]** | **[100 100]** | **[100 200]** |
|---|---|---|---|
| **CHLA9** | (43, 45) = 0.33 {tail-tail}<br>Num Fish = 15 (%15)<br>C = 0.43 ± 0.14 | (27, 29) =0.33 {tail-tail}<br>Num Fish = 15 (%15)<br>C = 0.33 ± 0.08 | (14, 15) = 0.45 {tail-tail}<br>Num Fish = 34 (%34)<br>C = 0.38 ± 0.03 |
| **CHLA10** | (46, 47) = 0.42{tail-tail}<br>Num Fish = 33 (%36)<br>(46, 83) = 0.51<br>{tail- branchial arches}<br>Num Fish = 39 (%42)<br>(47, 83) = 0.57<br>{tail- branchial arches}<br>Num Fish = 43 (%47)<br>C = 0.42 ± 0.13 | (30, 31) = 0.43{tail-tail}<br>Num Fish = 33 (%36)<br>(30, 35) = 0.51<br>{tail-branchial arches}<br>Num Fish = 40 (%43)<br>(31, 35) = 0.61<br>{tail- branchial arches}<br>Num Fish = 45 (%49)<br>C = 0.43 ± 0.03 | (15, 16) = 0.48{tail-tail}<br>Num Fish = 41 (%45)<br>(15, 18) = 0.69{<br>tail- branchial arches}<br>Num Fish = 62 (%67)<br>(16, 18) = 0.61<br>{tail- branchial arches}<br>Num Fish = 53 (%58)<br>C = 0.58 ± 0.03 |
| **CHLA258** | (38, 39) = 0.8 {VDA-VDA}<br>Num Fish = 4 (%4)<br>(39, 40) = 0.5 {VDA-VDA}<br>Num Fish = 4 (%4)<br>C = 0.55 ± 0.23 | (22, 23) = 0.36 {VDA-VDA}<br>Num Fish = 4 (%4)<br>(23, 24) = 0.44 {VDA-VDA}<br>Num Fish = 4 (%4)<br>C = 0.32 ± 0.09 | (11, 12) = 0.21 {VDA-VDA}<br>Num Fish = 5 (%5)<br>C = 0.31 ± 0.02 |
| **TC32** | (46, 83) = 0.56<br>{tail- branchial arches}<br>Num Fish = 44 (%44)<br>C = 0.40 ± 0.12 | (30, 35) = 0.58<br>{tail- branchial arches}<br>Num Fish = 45 (%45)<br>(31, 35) = 0.42<br>{tail- branchial arches}<br>Num Fish = 33 (%33)<br>C = 0.39 ± 0.03 | (15, 18) = 0.69<br>{tail- branchial arches}<br>Num Fish = 63 (%63)<br>C = 0.50 ± 0.03 |
| **TC32<br>shEF1CTR** | (45, 46) = 0.44{tail-tail}<br>Num Fish = 21(%23)<br>C = 0.50 ± 0.21 | (29,30) = 0.44{tail-tail}<br>Num Fish = 21 (%23)<br>C = 0.33 ± 0.04 | (14,15) = 0.47{tail-tail}<br>Num Fish = 27 (%30)<br>C = 0.39 ± 0.03 |
| **TC32<br>shEF1KD** | ([46,47], 83) = 0.54, 0.45<br>{tails- branchial arches}<br>Num Fish =41, 32 (%47, %37)<br>(36,46) = 0.53{1st ISV-tail}<br>Num Fish =32 (%37)<br>([11,27] 83) = 0.14, 0.34 {DLAV- branchial arches}<br>Num Fish =9, 24 (%10, %28)<br>C = 0.45 ± 0.17 | ([29,30,31], 35) = 0.47, 0.57, 0.46<br>{tails- branchial arches}<br>Num Fish = 33, 43, 33 (%38, %49, %38)<br>(20, 30) = 0.5{1st ISV-tail}<br>Num Fish = 32 (%37)<br>(11, 35) = 0.44<br>{DLAV- branchial arches}<br>Num Fish =32 (%37)<br>C = 0.37 ± 0.03 | (15, 18) = 0.68<br>{tails- branchial arches}<br>Num Fish =55 (%63)<br>(10, 15) = 0.55{1st ISV-tail}<br>Num Fish =42 (%48)<br>(6, 18) = 0.55<br>{DLAV- branchial arches}<br>Num Fish =41 (%47)<br>C = 0.49 ± 0.04 |

Under these criteria, CHLA9 exhibited robust coexistence among tail-region pairs at the 100 × 100 and 100 × 200 grids but not at the smaller 50 × 100 scale, indicating the presence of multiple colonies within the caudal hematopoietic tissue (CHT) rather than a single dominant colony in at least 15% of fish (Fig. 5c). In CHLA10, strong coexistence was observed between the tail and branchial arches across all grid sizes, occurring in more than 42% of the population. This suggests that for this cell line, these regions may share environmental characteristics fostering preferential localization of spreading cancer cells. Tail–tail coexistence pairs in CHLA10 were close to random expectation, although they occurred in 36% of fish, consistent with the presence of multiple tail colonies, similar to CHLA9. CHLA258 displayed weak coexistence patterns in the ventral wall of the dorsal aorta (VDA), indicating multiple colonies only in 4% of the population. TC32 exhibited patterns similar to CHLA10, with strong coexistence between the tail and branchial arches in at least 44% of fish. In contrast, TC32 shEF1 CTR showed weaker and less consistent tail coexistence across spatial scales. TC32 shEF1 KD demonstrated multiple robust coexistence pairs conserved across grid sizes, including tail–branchial arch pairs that exceeded the significance threshold and were present in at least 47% of the population. In TC32 shEF1 KD, weak coexistence was observed between the tail and first intersegmental vessel (ISV), likely due to the coarse definition of the ISV-containing box, which includes multiple anatomical regions and limits interpretability. Similarly, dorsal vessel–branchial arch coexistence appeared near the random threshold and was observed only at the 100 × 100 and 100 × 200 grids, but not at 50 × 100.

Overall, this analysis reveals non-random, condition-specific regional coupling of colonization patterns that are stable across spatial scales. This supports the notion that different cell lines harbor functional profiles, which specializes them for colony formation in select anatomical regions during organ-specific spreading. In particular, cell lines with higher spreading propensity such as CHLA10, TC32 and TC32 shEF1 KD, exhibit shared visiting coordinated regions of the branchial arches and tail regions, indicating these environments may be particularly conducive to cancer cell survival, proliferation, or retention.

## Discussion

In this study, we leverage the zebrafish embryo as an *in vivo* host to establish a systematic, and cost-effective platform for mapping the organotropic spreading of cancer cell types of different origin and molecular make-up. We extended a preliminary implementation of the Fish Metastatic Atlas (FMA)^27^ concept with novel technical solutions to animal alignment and aggregation, enabling robust statistical analysis of metastatic distributions across fish. We developed statistical metrics to quantify global and local spatial variation across distinct cell lines and applied this framework to compare spreading patterns in patient-matched primary and metastatic Ewing sarcoma tumors and cell lines, and under defined perturbations of the putative driver pathway.

The described FMA pipeline relies on a data-driven reference image rather than a rigid anatomical template, preserving inter-sample variability (e.g., variations in larval developmental stage, injection efficiency, imaging conditions) while minimizing distortion. We use the zebrafish vasculature as the registration channel because of its spatial richness, stereotypy during imaging, and direct relevance to cancer cell dissemination. The pipeline is modular and continuously expandable, allowing incorporation of new cell lines and imaging modalities without redesigning.

Cancer colonies are represented both as whole-image (whole organism) maps for global comparisons and as region-based frequency distributions to quantify preferential metastatic target areas. The region-based strategy supports anatomically defined regions and unbiased grid partitioning, enabling high-resolution analyses of specific organ–cancer interactions. Using these data, xenografting cancer cells into various organ- and tissue-specific zebrafish reporter lines could then validate the organ-specificity of metastasis. Furthermore, zooming into the identified metastatic regions of interest, such as the CHT, VDA or DLAV for various EwS cell lines, allows comparative studies of the interactions between cancer cells and the specific tumor microenvironments.

We analyzed interdependency across regions at the single-fish level. This systems-level approach frames metastasis as a networked disease rather than independent local events, revealing how colonization of one organ may associate with the probability of metastasis in another. Such analyses can be used to distinguish driver metastatic sites from endpoints, and uncover coordinated clonal behaviors across tissues, expanding the classic “seed and soil” concept into a multi-organ ecosystem. Coexistence of CHLA10, TC32 and TC32 shEF1 KD colonies in the distal tail (CHT, AVJ) and branchial arches is one interesting example. Both regions share distinctive vascular features: narrow, curved vessels with relatively slow blood flow, as seen in the dense vascular networks of the CHT/AVJ and the developing gill vasculature within the branchial arches. These mechanical and physiological conditions may promote stalling and enhance extravasation of circulating tumor cells, contributing to preferential colonization in both sites. Furthermore, these regions are also linked by function and developmental origin to major sites of EwS metastasis in humans - bone marrow and lungs, respectively. The CHT is an important hematopoietic niche in zebrafish larvae, supporting the development of hematopoietic stem cells (HSCs) between 2-5 dpf. In parallel, the thymus develops in the region of branchial arches and is colonized by HSCs from 3dpf onwards^46,47^. The brachial arches also support the development of the gills. Although bony fishes such as zebrafish lack lungs, gills and lungs share a common embryologic origin, and the epithelium of both organs derives from branchial arch endoderm^48^. Thus, the coexistence of cell colonization in the distal tail and branchial arch regions of the fish could also reflect microenvironmental conditions in these regions similar to those in common sites of metastasis in human Ewing sarcomas.

Looking forward, integration of the FMA with automated high-throughput imaging systems such as a vertebrate automated screening technology (VAST) bioimager system (Union Biometrica Inc.) will allow dynamic, longitudinal tracking of colonization events and rare processes such as extravasation at single-cell resolution. Future extensions could incorporate immune-competent contexts, multi-omic profiling, and perturbation screens to further dissect tumor–organ interactions. Ultimately, this framework provides a foundation for studying metastasis as a spatially structured, evolving, and system-wide disease, with implications for identifying targeted therapeutic strategies that could then further be studied in more time-consuming mammalian models.

## Acknowledgements

We would like to thank Stephan Daetwyler, Jungsik Noh, Felix Zhou, David Saucier, Reto Fiolka, Conor McFadden, and Hyeyun Christine Lee for fruitful discussions. This work was supported by National Institutes of Health grants K99CA283246 to H.M, U54CA268072 to G.D. and J.F.A funding from the Kate Amato Foundation to I.L. and the TJ Martell Foundation to J.F.A. J.F.A. is supported by the Alfred E. Mann Family Foundation Chair in Cancer Research. Software dissemination is funded by RM1 GM145399.

## Author Contributions

G.D. and J.A. set the overall goals of the project. H.M.-F. and G.D. conceived the technical concepts of the project. H.M.-F. and I.L. contributed to shaping the conceptual and biological framework of the study. H M.-F. performed all computational analyses and wrote the core code for the package, with contributions from Q.Z. I.L. and J.A. defined the experimental assays, and I.L. designed and performed all experiments. for the project. H.M.-F. wrote the first draft of the manuscript, and I.L., G.D., and J.A. contributed to writing and editing. All authors read and approved the final manuscript.

## Code Availability

This framework is open source and is available at https://github.com/DanuserLab/FishATLAS.

## Methods

### Zebrafish husbandry

*Danio rerio* were maintained at the AALAAC-accredited CHLA Research Aquatic Facility following established industry standards. The Tg(*kdlr:eGFP*) and Tg(*kdrl:dsRed*) fish were acquired from ZIRC Zebrafish International Resource Center (https://zebrafish.org).

#### Cell culture and cell line generation

All EwS cell-lines were cultured at 5% CO_2_ and 21% O_2_. CHLA9, CHLA10 and CHLA258 cells were cultured in IMDM (Gibco) supplemented with 20% fetal bovine serum (FBS), 1% antibiotic antimycotic and 1xInsulin-transferrin-Selenium (Gibco). TC32 were cultured using RPMI (Gibco) supplemented with 10% FBS and 1% antibiotic antimycotic. TC32 *EWSR1-FLI1* (EF1) shRNA cell line generated by the Grünewald lab^41^ was cultured in RPMI (Gibco) supplemented with 10% FBS and 1% antibiotic antimycotic. shRNA knockdown was induced by 1μg/mL doxycycline in cell culture for 72 hours prior to injection, and 10μg/mL doxycycline in E3 fish larval growth medium following injection until imaging. EF1 shRNA knockdown was verified by western blot using rabbit mAb to Fli1 (Abcam ab133485) (Supplementary Fig 1a). Fluorescent constructs were introduced into cells using ViraPower Lentiviral Packaging Mix (Invitrogen, Cat #K497500) and Lipofectamine 3000^TM^ Transfection reagent (Invitrogen, Cat #L3000015). The F-Tractin GFP and F-Tractin mRuby2 constructs contain residues 9–52 of the enzyme IPTKA30 fused to eGFP or mRuby2, respectively^21^.

#### Ewing sarcoma xenografts in zebrafish

Xenografts into the ventral perivitelline space (PVS) of zebrafish larvae were performed as previously established by the Fior lab^37^ with minor modifications. Briefly, zebrafish embryos were collected in E3 embryo medium and treated with 0.1mM phenylthiourea (PTU) starting at 1 day post fertilization (dpf) to prevent pigmentation. At 2dpf, larvae were anesthetized using Tricaine methanesulfonate (Sigma) and aligned on an agarose plate for injection. Cells were prepared in culture medium at a concentration of 1x10^5^ cells/μL and kept on ice. Injection needles (Sutter Instrument borosilicate glass O.D 1.0, I.D 0.75, Cat# B100-75-10) were pulled with Sutter Instrument MODEL P-2000 (program: Heat 300, Fil 4, Vel 30, Del 100, Pil 75). Approximately 500 cells were injected into the PVS through the yolk using Warner Instruments pico-liter injector (PLI-90A) with an injection time of 0.1sec and pressure meter on 0.5-1.2 (depending on the needle). Successful injections (visible cell mass in the PVS) were raised at 34°C in PTU-treated E3 embryo medium for three days before imaging to allow time for stabile metastatic colonies to form. At 3 days post injection (dpi), xenografted larvae were anesthetized with Tricaine, screened for a visible PVS cell mass and mounted in 1% low gelling temperature agarose (Sigma Cat# A9414) supplemented with Tricaine. Accurate aligning of the larvae (flat against the plate with no tilting) while mounting was ensured for best results during downstream registration during image analysis. The larvae were imaged using a Leica M205FA stereoscope for brightfield, vasculature, and cancer cell channel.

#### Image preprocessing and data-driven reference image

A custom-package in MATLAB was developed to analyze the images. We used all the registration using vasculature structures images. Prior to registration, all images in both channels were positioned on the fish body at the center with the eye positioned on the left and the dorsal side facing upward. A data-driven reference image can be generated before the analysis for a single population or across multiple populations in a separate module using a groupwise-based deformation of all fish images from several conditions. In the groupwise approach, there is no final reference image across the population, and all images are simultaneously registered together in a population to eliminate the biased associated to a single image. Each image is registered as a new image with a displacement field that minimizes the intensity variance under the constraint that the average deformation over images is zero. We chose the median of all registered images as a reference image for further analysis to evaluate organ-specific patterns of cancer cells across conditions. Schematics were created with BioRender.com.

#### Image registration

Depending on reference availability, fish were aligned using either pairwise or groupwise registration strategies. Both strategies include rigid geometric transformation that includes translocation, rotation and scaling, and deformable transformations either using isotropic total variation of regularization of displacement (*imregdeform* function in MATLAB) or an intensity-based metric using diffeomorphic demons algorithm (i*mregdemons* function in MATLAB). When a reference fish was available, either user-selected image or pre-calculated data-driven reference image, each fish aligned with the reference image in a pairwise strategy. In this case, the reference image is similar across all conditions, and comparing analysis across the conditions can be easily done. In the absence of a reference image, fish were rigidly aligned to a randomly selected image from the dataset, followed by a groupwise-based deformation process. Users can also select an image from the dataset instead of relying on a random choice. We generated a semi-reference image within each population as the average of registered images after groupwise deformation. For statistical analysis across conditions, these reference images were realigned to each other to define a global reference image across all conditions. For both rigid geometric transformation and deformable transformation, we applied a filtering approach to discard the not-well aligned registered fish using the correlation coefficient between the registered image and data-driven reference image.

For both rigid and deformable transformations, a filtering step was applied to discard poorly aligned fish based on the correlation coefficient between the registered image and the corresponding data-driven reference image. Correlation thresholds were set to 0.70 for rigid registration and 0.82 for deformable registration. In the groupwise registration approach, the groupwise deformation step was repeated after discarding poorly aligned images until convergence was achieved, defined as a correlation coefficient of at least 0.82 or a maximum of five iterations.

#### Validation of image registration

To validate the accuracy of image registration, we developed a user-friendly interface to manually annotate biological landmarks and anatomical sites on the vasculature images of individual fish. For each experimental condition, ten fish were randomly selected, and 14 anatomical landmarks were manually marked on each fish (Supplementary Fig 3a). To reduce bias associated with manual annotation, each landmark was selected three times per fish, and the average coordinate across the three selections was used for subsequent analysis.

The same anatomical landmarks were manually annotated on the reference image. Using the geometric transformations and displacement fields obtained from the registration process, the averaged landmark coordinates from each fish were mapped onto the reference image. For each landmark, the Euclidean distance between the transformed coordinates from the individual fish and the corresponding reference landmark was computed. Registration precision for each anatomical site was summarized using boxplots across the ten fish, providing a quantitative assessment of registration accuracy at individual anatomical locations.

#### Cancer colony detections: binary and point representations and colony

Cancer colonies were first identified by binarizing xenografted cell images using Otsu’s multilevel thresholding (two threshold levels). Prior to thresholding, each image was smoothed using a Gaussian low-pass filter to suppress high-frequency background noise and random intensity fluctuations, thereby improving the robustness of the segmentation. The Gaussian filter performs a weighted averaging of neighboring pixels based on a Gaussian kernel, reducing noise while preserving the overall morphology of the cancer colonies. The resulting binary images were mapped onto the reference image obtained from the vasculature-guided registration process using the corresponding geometric transformations and displacement fields. Accumulation of the mapped binary images onto the reference image represents organ-specific patterns for each cell line.

To quantify site-visitation frequency for each cell line, cancer colonies were additionally detected as point sources corresponding to local intensity maxima fitted with a Gaussian model (σ = 6, α = 0.0005) using a previously published detection package^49^. Dimmer maxima related to non-specific cancer colonies were removed by applying an intensity threshold of 4% of the maximum signal, which effectively filtered false detections. The final cancer colony coordinates were then geometrically transformed into a corresponding final reference image using the rigid transformation and deformation fields from the registration process. For each experimental condition and cell line, point coordinates from individual fish were accumulated into the final reference image.

In both binary and point-based analyses, a binary mask of the fish body, generated from the vasculature image, was used to exclude pixels, or detected maxima located outside the fish body.

#### Comparison of organ-specific patterns in mapped binary images

Pairwise comparisons of organ-specific cancer hotspot patterns across experimental conditions were performed using the Jaccard similarity coefficient with permutation testing. Accumulated hotspot maps were first normalized by the number of fish per condition to enable fair comparisons across conditions. The resulting accumulation maps are grayscale, reflecting the fraction of fish contributing to each spatial location. To compute the Jaccard index, the normalized accumulation maps were converted to binary images using a user-defined visitation probability threshold. The Jaccard index was then calculated using the standard formulation:

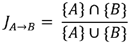

where {A} and {B} denote the sets of pixels in the accumulators of condition A and condition B, respectively, whose values are above a user-defined visitation probability threshold. We chose a 1% threshold, meaning that if a hotspot existed in even one fish, that region was retained. To generate a similarity matrix across all condition pairs, we used a bootstrap-based permutation test to calculate permutation p-values.

In addition to the standard Jaccard similarity, we computed a weighted Jaccard index using the normalized grayscale accumulation maps without applying thresholding. The weighted Jaccard index was defined as:

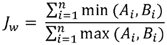

where and are the normalized accumulation values at the th pixel for conditions A and B, respectively, and is the total number of pixels. These metrics capture pixel-wise similarity based on the actual contribution of each condition at each spatial location. Likewise, we computed the similarity matrix using the permutation-based p-value using a bootstrapping approach for all pairwise comparisons.

#### Spatial analysis of organ-specific cancer colony visitation patterns

Using accumulators of the point patterns, we compared region visitation frequency per experimental condition. Number of nearest neighbors within a user-defined radius (R = 40 pixels, corresponding to the distance between two nearby vessels in IDV) was computed for each point. To correct for edge effects that underestimate the number of neighbors near the fish boundary, we used a fish mask created by manually tracing the edge of the reference image. For each point, we calculated the fraction of the circle (centered at that point with radius 40 pixels) lying outside the fish body and multiplied the number of neighbors by this fraction to correct for underestimation. Points were then clustered and labeled based on the maximum number of nearest neighbors within this radius, using arbitrary population thresholds (N = 1, 10, 50, 100, 150). A cluster population matrix was generated for each experimental condition and normalized by the number of fish to account for variability in sample size. This analysis was also used to estimate the minimum number of fish required to achieve stable spatial patterns within a single condition.

Top-ranked regions in the vasculature channel of the computed reference image were selected by considering an R = 40-pixel neighborhood around points with the highest population density (top two regions per experimental condition).

To assess whether point patterns were clustered or isolated without relying on predefined population thresholds, observed point patterns were compared against random distributions. For each condition, the same total number of points was randomly distributed within the fish body mask. The number of nearest neighbors within R = 40 pixels was computed for each random realization. This process was repeated 1,000 times to generate a reference distribution of nearest-neighbor counts. Observed point patterns were then compared to the random distributions, and clustered versus isolated points were identified as those falling within the left or right tails of the random distribution histogram. This approach enabled calculation of p-values and identification of statistically clustered or isolated regions, independent of the choice of high versus low population thresholds, which may vary across experimental conditions.

#### Spatial enrichment of cancer colonies

We used an annotation platform to define user-selected anatomical regions for comparing experimental conditions within predefined spatial domains. Binary masks were generated for each annotated region, and visitation frequencies were quantified by counting the number of cancer colony visits within each region across fish. The average distribution of number of colonies is visualized in a bar graph for each experimental condition.

We calculated the p-value of spatial enrichment based on the distribution of cancer colony counts within each region for a given condition. To establish a null distribution, we generated 1,000 randomly distributed patterns across the fish body for that condition and counted the number of events within the region for each pattern. The p-value is defined as the fraction of these random patterns that exhibit a higher number of events than the observed pattern.

#### Region-based analysis

The position of the fish body in each image was identified using a mask generated from the vasculature images. On the reference image, a series of boxes with user-defined sizes were overlaid across the fish body. For each box, the number of cancer colony points was counted, producing a matrix of population densities that represents spatial distribution across the fish body. Experimental conditions were compared using these defined regions, and population counts were normalized to the number of fish per condition.

For pairwise comparisons, we calculated the ratio of dominant cell lines or experimental conditions per region using the normalized population values. Statistical significance of dominance was assessed using the Wisconsin rank test.

#### Interdependency analysis of regions for cancer colonies

For each experimental condition, the same box grid was used to measure the number of points per box, representing the visitation frequency at fish level. A matrix of events per fish was generated for the box grid (Figure SI8). Coexistence between regions (boxes) at fish level was calculated for each pair of (box1, box2) using Jaccard index that is a ratio of number of fish that has at least one point at box 1 and box 2, divided by the number of fish that has at least one point in either box1 or box2. A threshold of 0.4 was applied to identify highly correlated regions. Correlation and anti-correlation patterns between regions were visualized using a chord diagram, providing insight into the interdependency of spatial regions for cancer colony distribution.

---

**Supplementary Figure 1.**
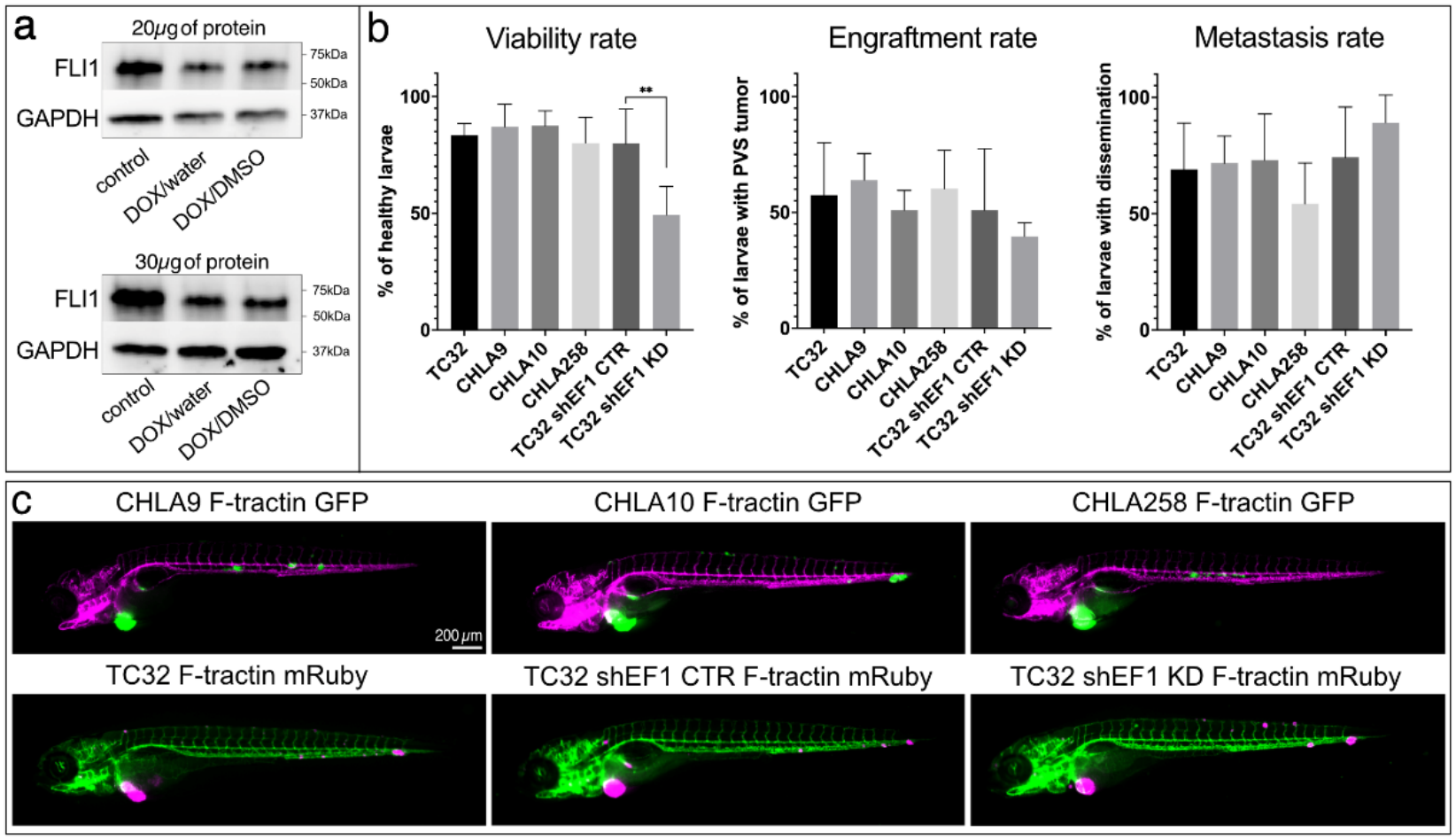
Xenograft Performance Across EwS Cell Lines. a) Two separate Western blots to validate EWSR1::FLI1 (EF1) downregulation upon doxycycline treatment of TC32 shEF1 cell line. Cells were treated for 72h with water- or DMSO-soluble doxycycline and analyzed with FLI1 antibody for EF1 fusion protein (70kDa) together with GAPDH control (37kDa). Both water- and DMSO-soluble doxycycline treatment led to ∼50% downregulation of EF1 (quantification not shown). b) Xenografting efficiency is similar for all studied EwS cell lines. The viability rate (% of healthy larvae at 3dpi) is significantly lower only in doxycycline-treated TC32 shEF1 KD line due to toxicity of the drug. The engraftment rate (% of healthy larvae with a PVS tumor at 3dpi) and metastasis rate (% of successfully engrafted larvae with tumor cells also outside of the PVS at 3dpi) did not vary considerably between the cell lines studied (ANOVA, **p<0.005). c) Representative images of 3dpi xenografted fish for all EwS cell lines studied. CHLA9, CHLA10 and CHLA258 cells were labelled for F-tractin GFP and xenografted into Tg(*kdrl:dsRed*) fish whereas TC32 cells were labelled for F-tractin mRuby and xenografted into Tg(*kdrl:eGFP*) fish.

**Supplementary Figure 2.**
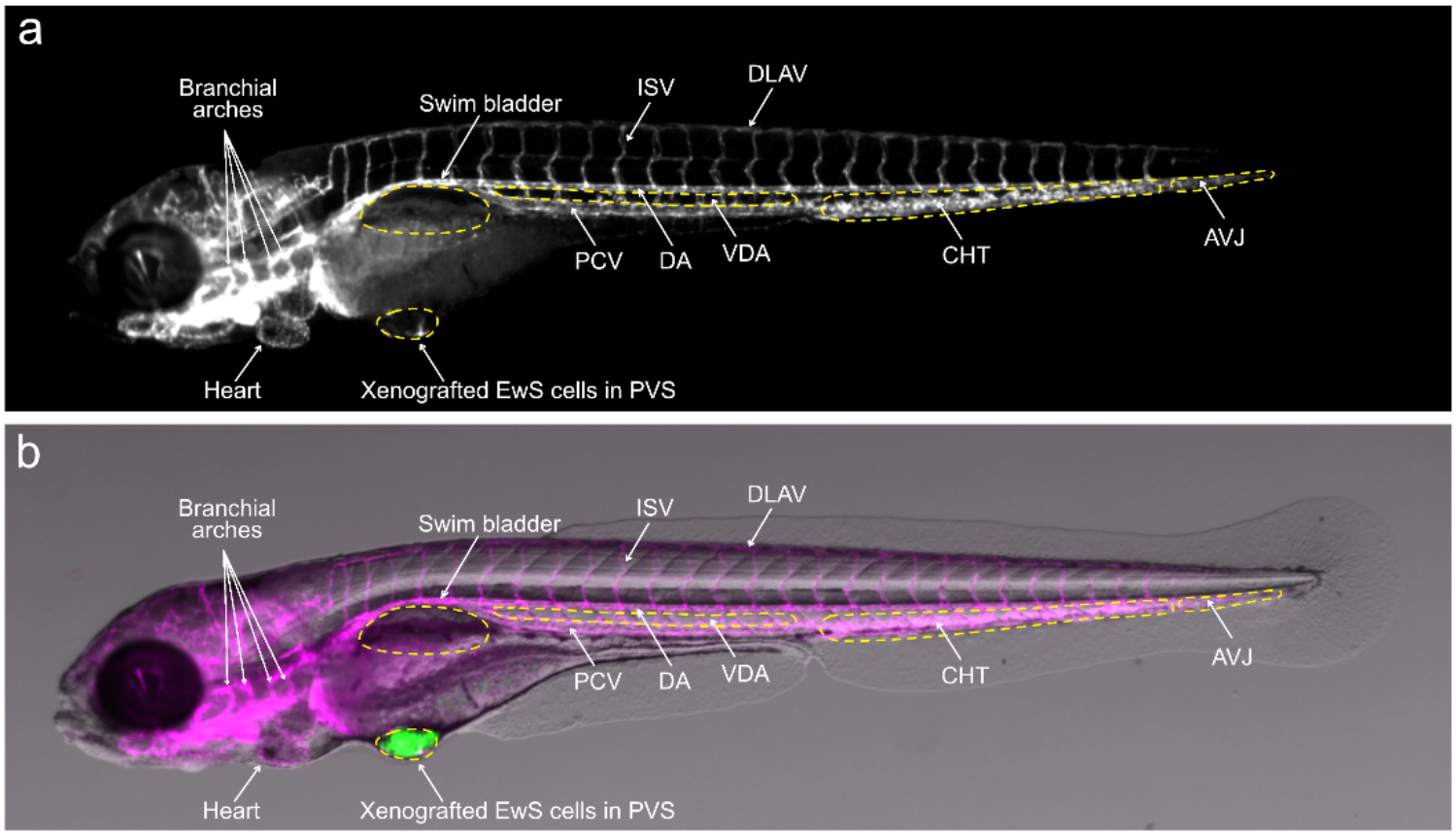
Fish Anatomy. Annotated zebrafish larvae at 5 days post fertilization (dpf)/3 days post injection (dpi) showing Tg(kdrl:dsRed) blood vessels (a) or blood vessels together with a brightfield anatomy image (b). Note the xenografting site (green EwS cells in the PVS) and the main regions of cell accumulation (the hematopoietic stem cell niche CHT and the tail AVJ). CHLA10 and TC32 EF1 knock down cells often also migrate to midline (ISV) and dorsal vessels (DLAV), whereas CHLA9 and CHLA258 cells accumulate in the VDA, which at the time of xenografting (2dpf) consists of a hematopoietic stem cell niche (aorta-gonad-mesonephros, not shown here). Abbreviations: AVJ - arteriovenous junction, CHT - caudal hematopoietic tissue, DA - dorsal aorta, DLAV - dorsal longitudinal anastomotic vessel, EwS -Ewing sarcoma, ISV - intersegmental vessel, PCV - posterior cardinal vein, PVS – perivitelline space, VDA – ventral wall of the dorsal aorta.

**Supplementary Figure 3.**
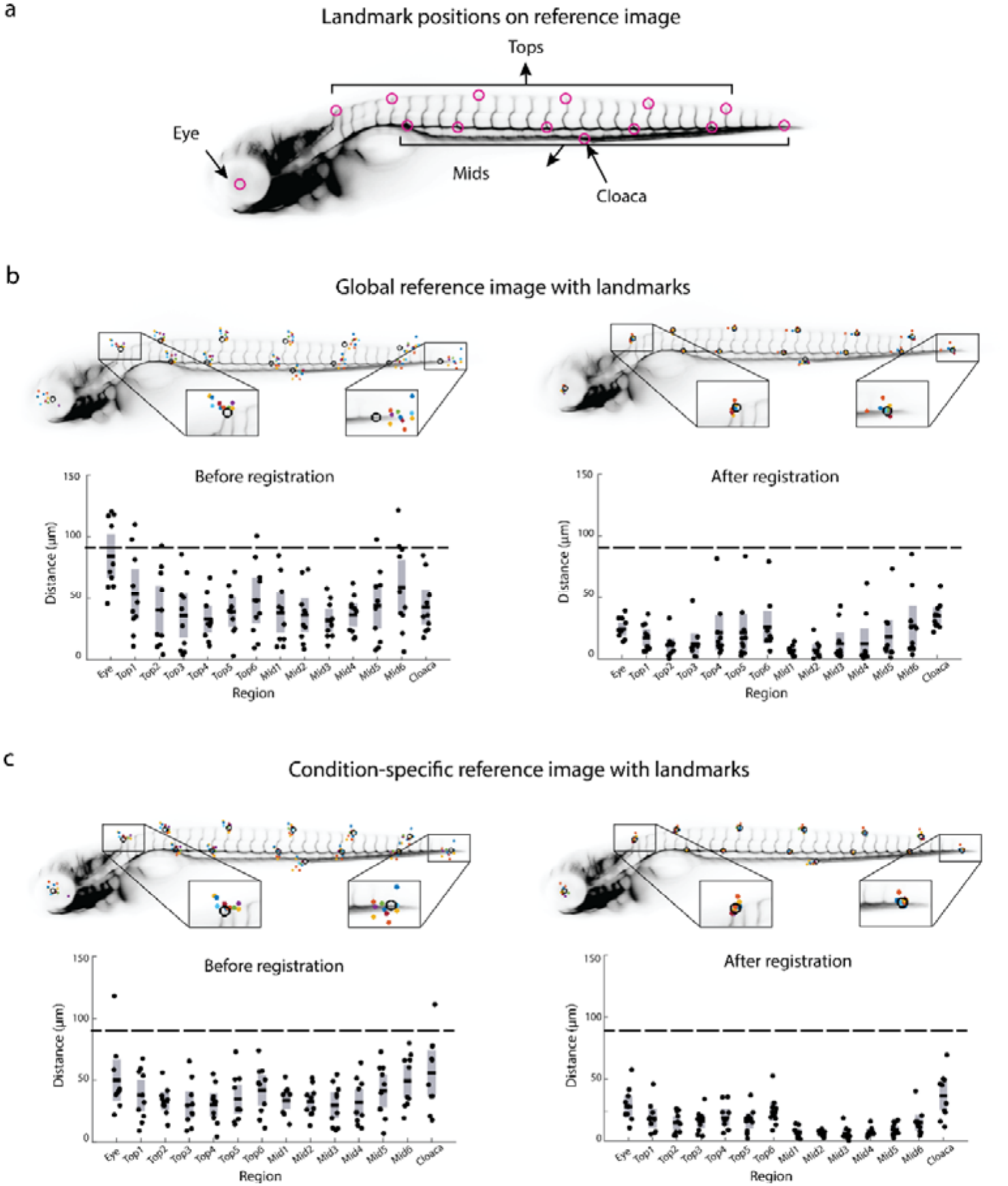
Validation of Vasculature-Based Registration Using Landmarks. a) Positions of fourteen landmarks on the vasculature reference image. One landmark is placed at the eye; six landmarks are positioned along the dorsal longitudinal anastomotic vessel (DLAV), starting at the first intersegmental vessel (ISV) and spaced by five ISVs between consecutive landmarks toward the tail; six landmarks are placed along the midline horizontal vessels, starting from the junction point, with three ISVs between the first and second landmarks and five ISVs between subsequent landmarks; the final landmark is positioned at the cloaca. b) Fourteen landmarks (color-coded) are shown for ten randomly selected CHLA9-injected fish, with black circles indicating landmark positions on the reference image. A global reference image was generated across all six conditions, with a common fish template applied prior to condition-specific analysis. Landmark positions are shown before (left) and after (right) registration. Corresponding boxplots summarize distances between landmarks from the ten fish and the reference image, with individual distances overlaid as scatter points. The dashed line represents the distance between adjacent intersegmental vessels (ISVs). c) Same as (b), except that the reference image was generated separately for each condition when a fish reference was unavailable.

**Supplementary Figure 4.**
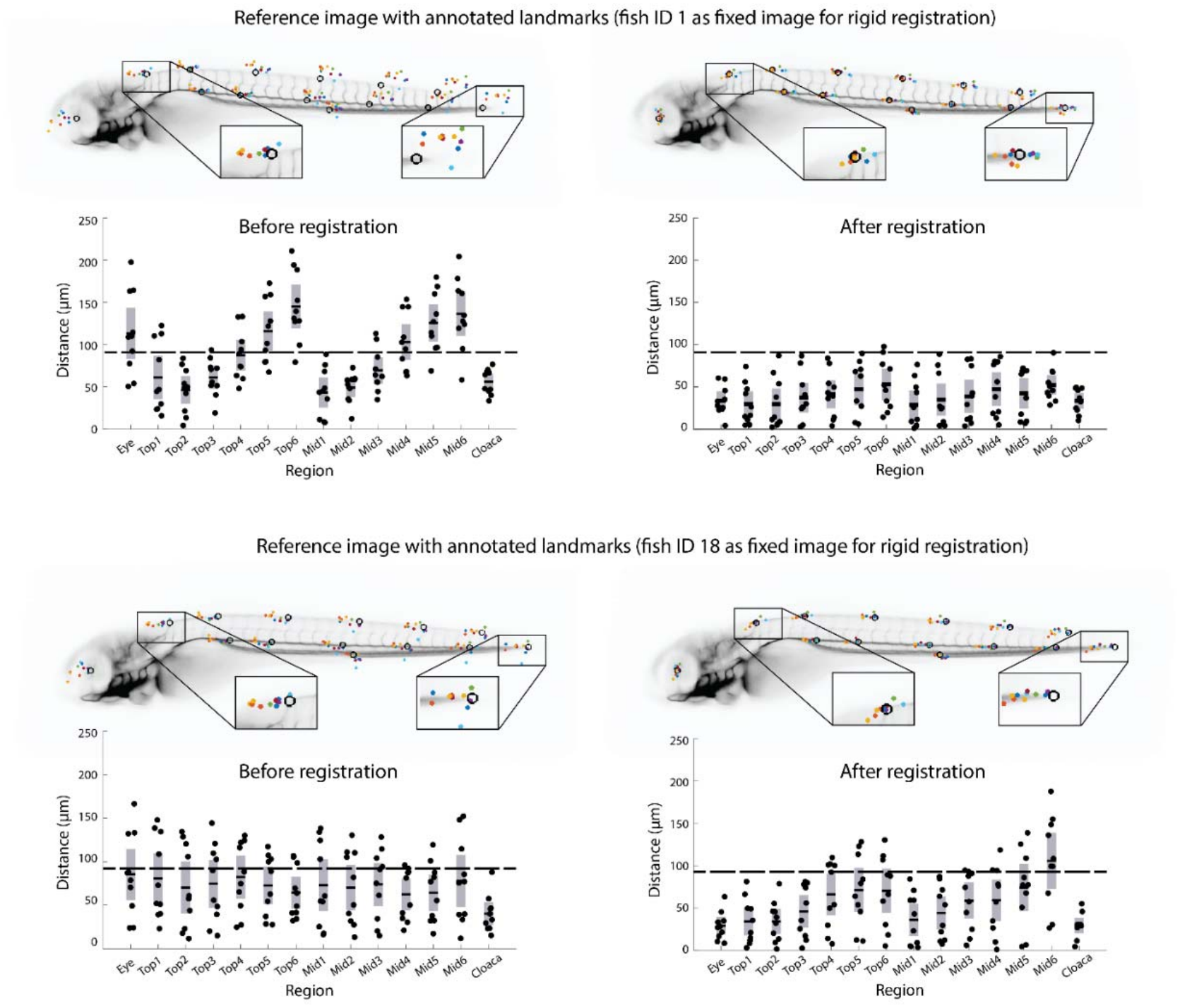
Validation of Rigid Registration with Fixed Reference Fish Using Landmarks. Fourteen landmarks (color-coded) are shown for ten randomly selected individual TC32-injected fish, with black circles indicating landmark positions on the reference image. The Fish ID 1 (top) and Fish ID 18 (bottom) were used as fixed images for rigid registration. Boxplots summarize distances between landmarks on the ten fish and the final reference image, with individual distances overlaid as scatter points.

**Supplementary Figure 5.**
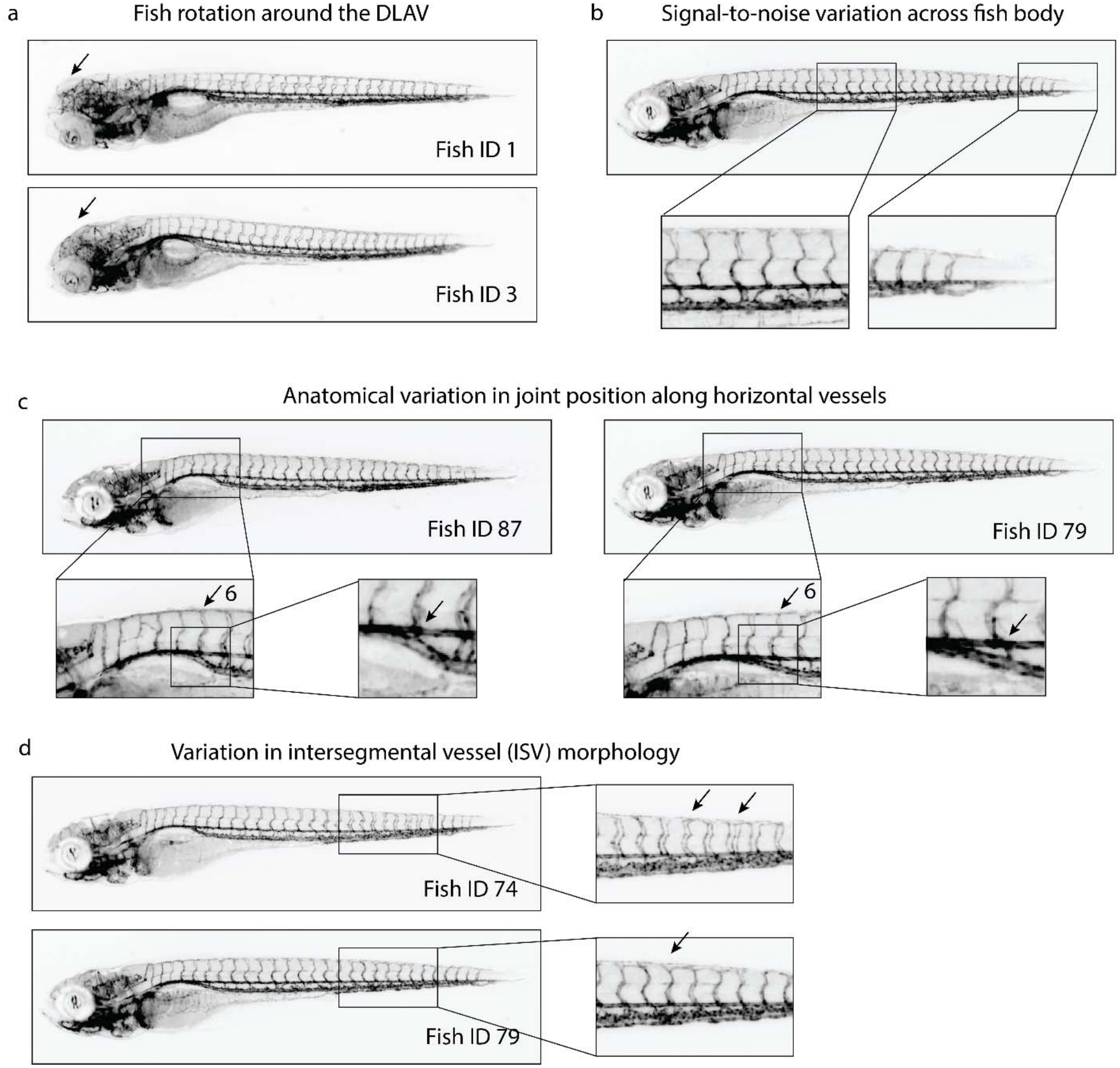
Sources of Anatomical and Imaging Variability in the Fish Dataset. a) Fish rotation around the dorsal longitudinal anastomotic vessel (DLAV) for two examples in the TC32 dataset: Fish ID 1 and Fish ID 3. b) Image signal variation across the fish body, with a higher signal-to-noise ratio in the middle region compared to the tail. c) Anatomical variation in the dataset, illustrated with two examples in TC32-injected fish image: Fish ID 87 has the joint point in the middle horizontal vessel at the 6th intersegmental vessel (ISV), whereas Fish ID 79 has the joint point between the 5th and 6th ISVs. d) Variation in intersegmental vessels (ISVs). Individual ISVs can contain one, two, or three parallel vessel lines, depending on the fish and region.

**Supplementary Figure 6.**
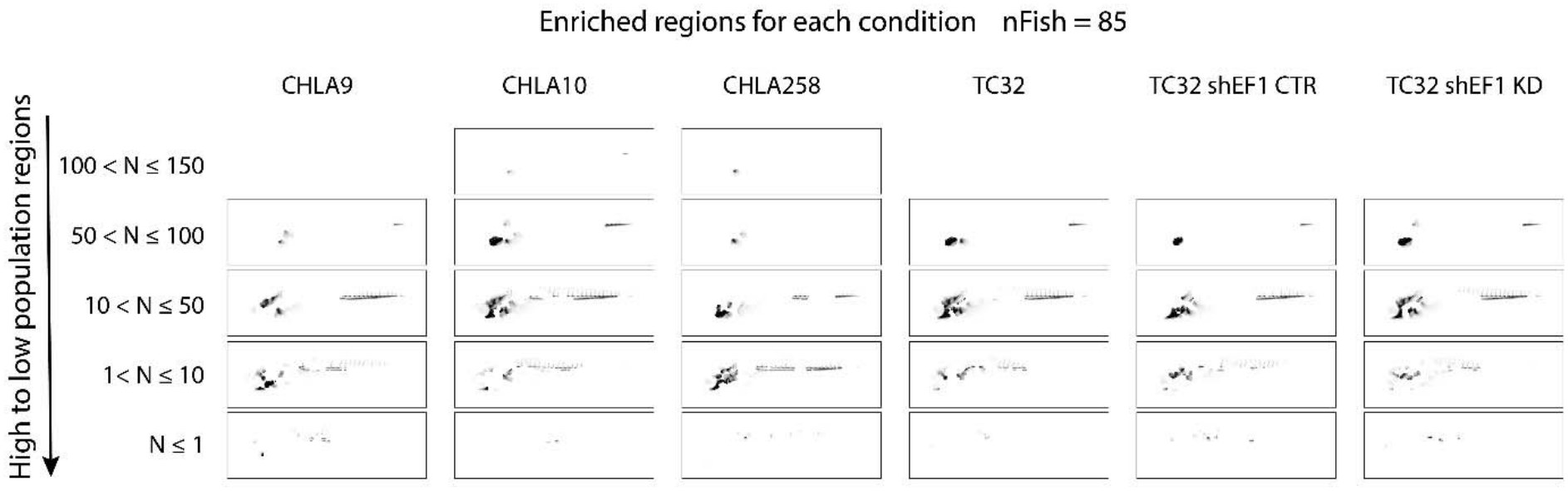
Regional Mapping by Local Cell Density. Regions of the fish body on the vasculature reference image based on local population density. Regions are masked according to the number of nearest neighbors within a radius of r = 40 pixels, using five bins corresponding to maximum neighbor counts of 1, 10, 50, 100, and 150. Regions are ranked from highest to lowest local population density and are shown for six experimental conditions: CHLA9, CHLA10, CHLA258, TC32, TC32 shEF1 Control, and TC32 shEF1 KD.

**Supplementary Figure 7.**
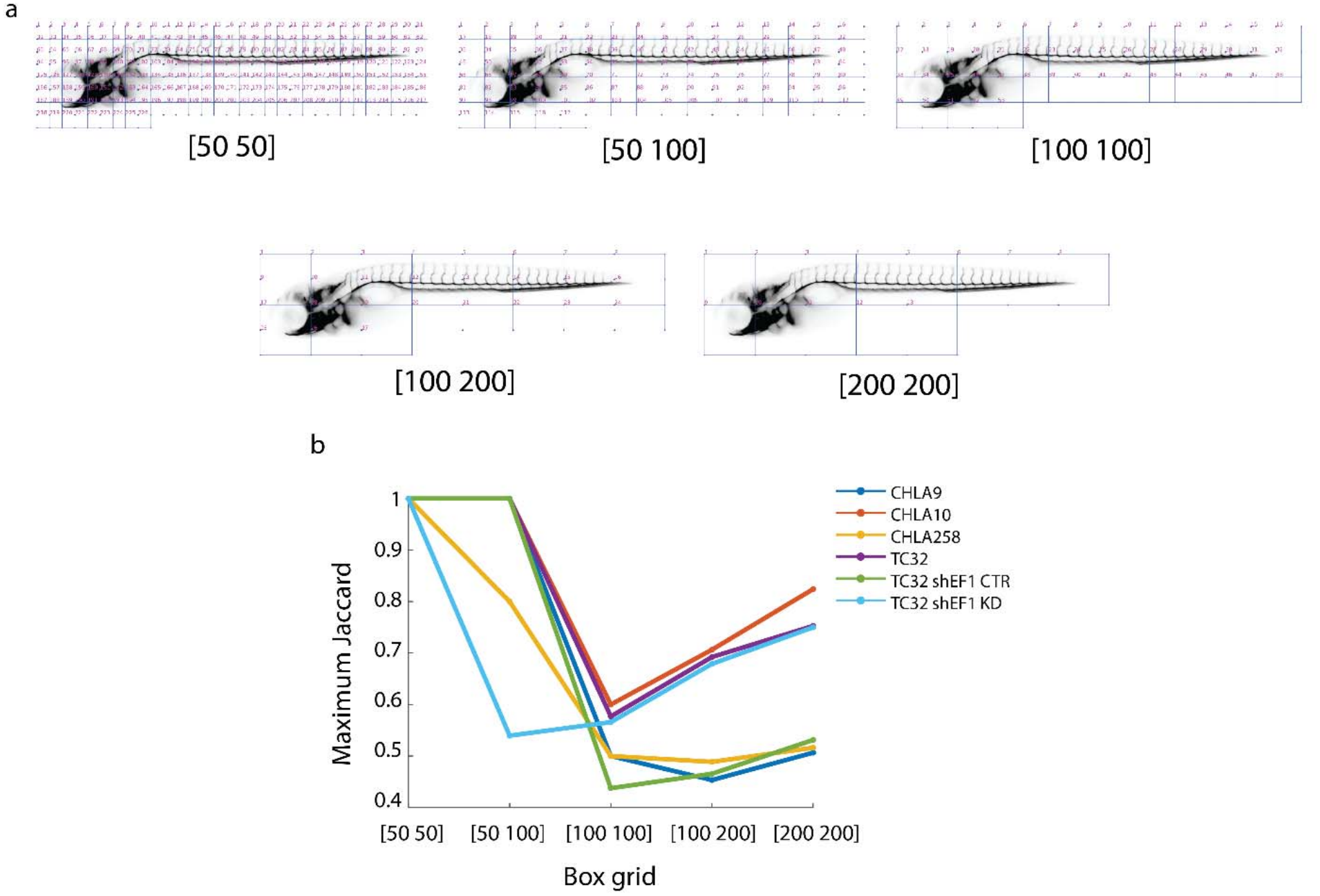
Box Grid–Based Analysis of Cancer Colony. a) Box grids overlaid on the reference image with box sizes of [50 × 50], [50 ×100], [100 × 100], [100 × 200], and [200 × 200] pixels. b) Maximum co-occurrence values across five box sizes are shown for all six conditions. Each point represents the maximum co-occurrence value across all box pairs for a given box size, defined as the ratio of observing at least one cancer colony in both boxes of a pair to observing a colony in either box.

**Supplementary Figure 8.**
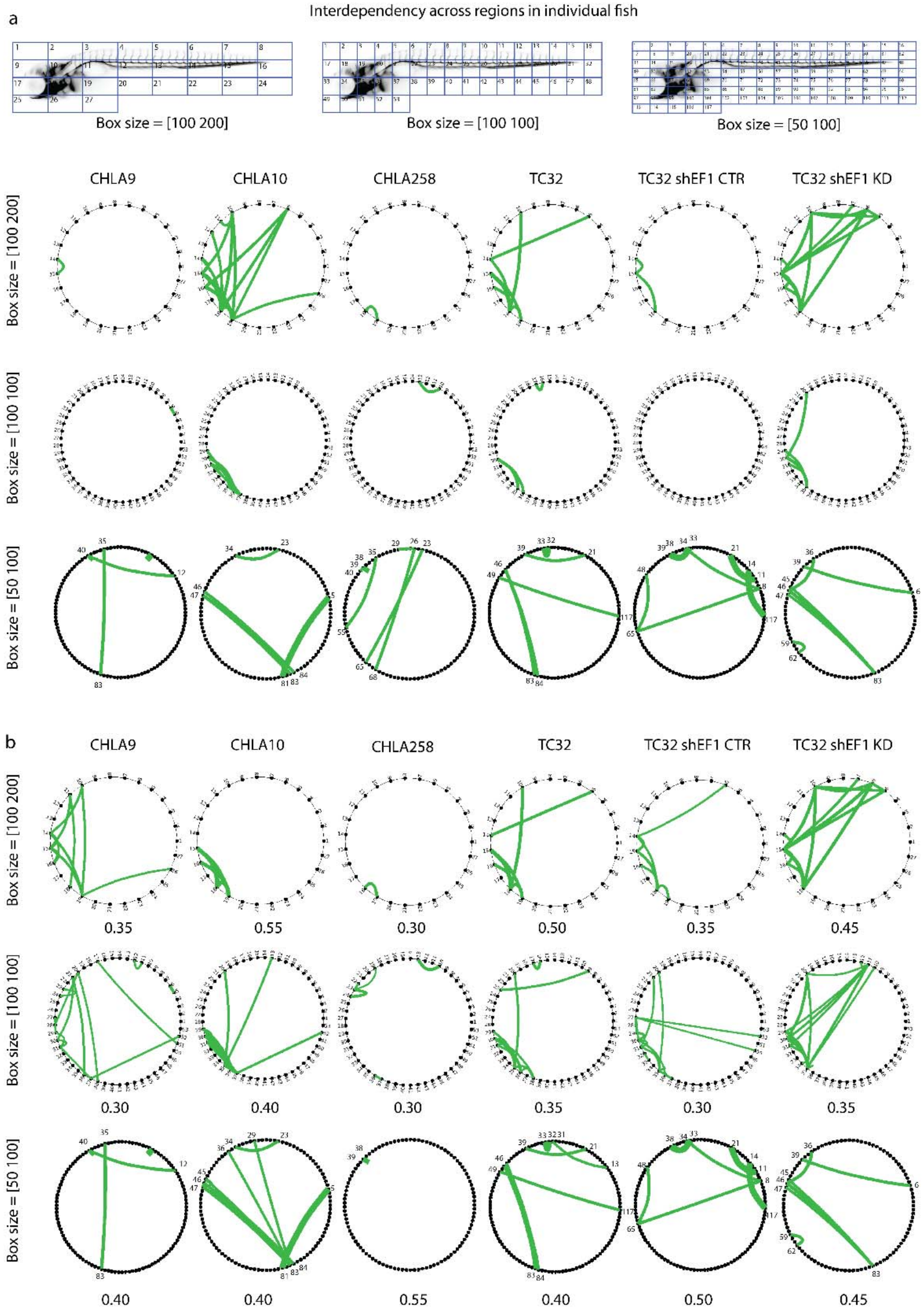
Region Interdependency Across Grid Sizes. a) Box grids overlaid on the reference image with box sizes of 100 × 200, 100 × 100, and 50 × 100 pixels. b) Region interdependency for each experimental condition. Chord diagrams depict region pairs with high absolute co-existence ratios, quantified using the Jaccard index (threshold ≥ 0.45). c) Chord diagrams of co-occurrence across three grid sizes, similar to (b), using thresholds determined for each condition and grid size based on a random distribution.

**Supplementary Figure 9.**
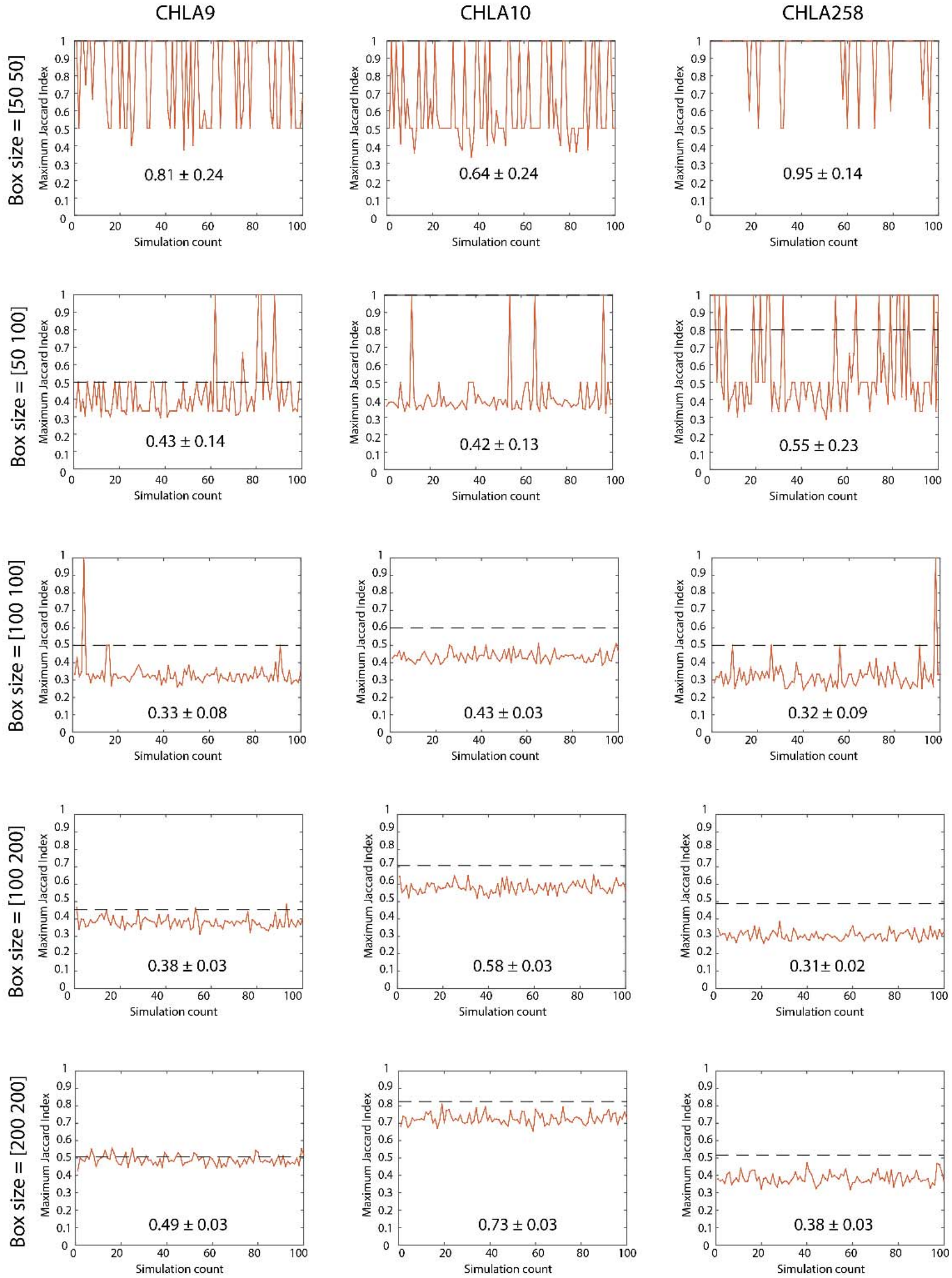
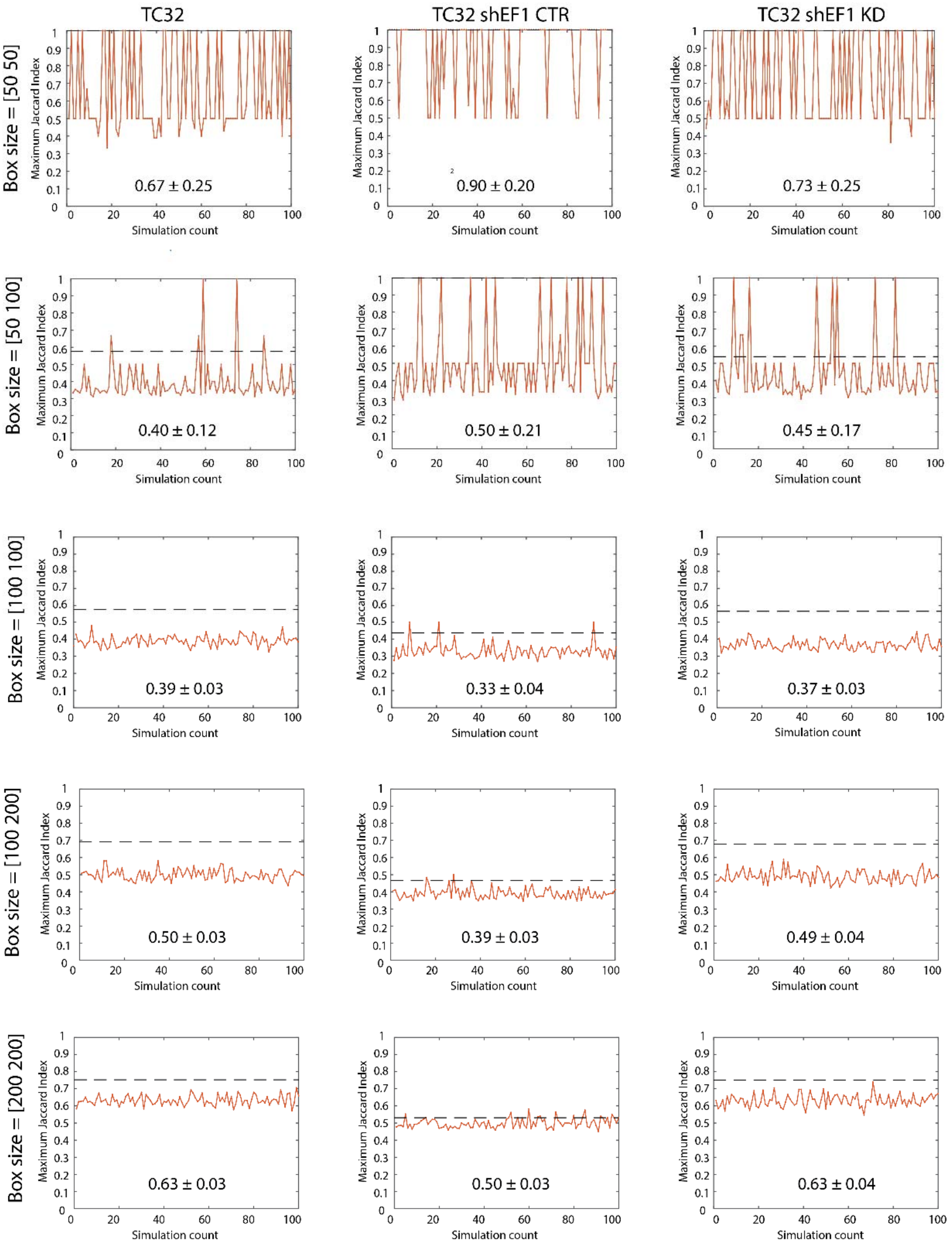
Significance Thresholds for Co-occurrence from Random Distributions. Maximum co-occurrence values across five box sizes are shown for each condition. Each subplot displays the maximum co-occurrence value across all box pairs for a given condition and box size. A total of n = 100 random distributions were generated for each condition and box size, and the resulting maximum co-occurrence values are summarized in each plot, with the mean and standard deviation indicated. The dashed line represents the maximum co-occurrence value of the original point pattern across the corresponding box grid.

**Movie1-6. Visualization of Cancer Colony Registration and Representation** Cancer colony registration and mapping for six experimental conditions: CHLA9, CHLA10, CHLA258, TC32, TC32 shEF1 control, and TC32 shEF1 knockdown. Each movie shows vasculature (purple) and cancer colonies (cyan) for individual fish, including raw images (top), registered binary colony masks on the reference vasculature (middle), and the reference image with colonies represented as red points (bottom).

